# Non-linear Auto-Regressive Models for Cross-Frequency Coupling in Neural Time Series

**DOI:** 10.1101/159731

**Authors:** Tom Dupré la Tour, Lucille Tallot, Laetitia Grabot, Valérie Doyère, Virginie van Wassenhove, Yves Grenier, Alexandre Gramfort

## Abstract

We address the issue of reliably detecting and quantifying cross-frequency coupling (CFC) in neural time series. Based on non-linear auto-regressive models, the proposed method provides a generative and parametric model of the time-varying spectral content of the signals. As this method models the entire spectrum simultaneously, it avoids the pitfalls related to incorrect filtering or the use of the Hilbert transform on wide-band signals. As the model is probabilistic, it also provides a score of the model “goodness of fit” via the likelihood, enabling easy and legitimate model selection and parameter comparison; this data-driven feature is unique to our model-based approach. Using three datasets obtained with invasive neurophysiological recordings in humans and rodents, we demonstrate that these models are able to replicate previous results obtained with other metrics, but also reveal new insights such as the influence of the amplitude of the slow oscillation. Using simulations, we demonstrate that our parametric method can reveal neural couplings with shorter signals than non-parametric methods. We also show how the likelihood can be used to find optimal filtering parameters, suggesting new properties on the spectrum of the driving signal, but also to estimate the optimal delay between the coupled signals, enabling a directionality estimation in the coupling.

**Author Summary:** Neural oscillations synchronize information across brain areas at various anatomical and temporal scales. Of particular relevance, slow fluctuations of brain activity have been shown to affect high frequency neural activity, by regulating the excitability level of neural populations. Such cross-frequency-coupling can take several forms. In the most frequently observed type, the power of high frequency activity is time-locked to a specific phase of slow frequency oscillations, yielding phase-amplitude-coupling (PAC). Even when readily observed in neural recordings, such non-linear coupling is particularly challenging to formally characterize. Typically, neuroscientists use band-pass filtering and Hilbert transforms with ad-hoc correlations. Here, we explicitly address current limitations and propose an alternative probabilistic signal modeling approach, for which statistical inference is fast and well-posed. To statistically model PAC, we propose to use non-linear auto-regressive models which estimate the spectral modulation of a signal conditionally to a driving signal. This conditional spectral analysis enables easy model selection and clear hypothesis-testing by using the likelihood of a given model. We demonstrate the advantage of the model-based approach on three datasets acquired in rats and in humans. We further provide novel neuroscientific insights on previously reported PAC phenomena, capturing two mechanisms in PAC: influence of amplitude and directionality estimation.

## Introduction

The characterization of neural oscillations, which are observed in the mammalian brain at different temporal and spatial scales, have given rise to important mechanistic hypotheses regarding their functional role in neurosciences (e.g. [Buzs´aki, 2006, Fries, 2015]). One working hypothesis suggests that the coupling across neural oscillations may regulate and synchronize multi-scale communication of neural information within and across neural ensembles [Buzsáki, 2010, Fries, 2015]. The coupling across different oscillatory activity is generically called cross-frequency-coupling (CFC) and has started to receive much attention [Jensen and Colgin, 2007,Lisman and Jensen, 2013,Canolty et al., 2006,Canolty and Knight, 2010,Hyafil et al., 2015]. The most frequent instance of CFC consists in the observation that the power of high frequency activity is modulated by fluctuations of low-frequency oscillations, resulting in phase-amplitude-coupling (PAC). Other instances of CFC include phase-phase coupling [Tort et al., 2007, Malerba and Kopell, 2013], amplitude-amplitude coupling [Bruns and Eckhorn, 2004, Shirvalkar et al., 2010], and phase-frequency coupling [Jensen and Colgin, 2007, Hyafil et al., 2015]. By far, PAC is the most reported CFC in the literature.

Seminally, PAC was described in local field potential (LFP) of rodents displaying a modulation of gamma band power (40–100 Hz) as a function of the phase of their hippocampal theta band (5–10 Hz) [Bragin et al., 1995, Tort et al., 2008]. Parallel recordings in different brain areas in behaving rodents have also highlighted differences in PAC between brain areas (e.g., hippocampus and striatum) at specific moments during a goal-oriented behavior, both in terms of which high-frequency range and how narrow-band the low frequency is [Tort et al., 2008]. PAC may promote cellular plasticity underlying memory formation [Axmacher et al., 2006]. In humans, theta (4–8 Hz)/gamma (80–150 Hz) PAC was described in human auditory cortex during speech perception [Canolty et al., 2006]. In more recent work, theta/gamma PAC was reported during the processing of auditory sequences in both humans and monkeys [Kikuchi et al., 2017], during working memory maintenance in human hippocampus [Axmacher et al., 2010], and during serial memory recall using non-invasive human magnetoencephalography (MEG) [Heusser et al., 2016]. PAC has been proposed to support the maintenance of information and to play an important role in long distance communication between different neural populations, considering that slow oscillations can propagate at larger scales than fast ones [Jensen and Colgin, 2007, Khan et al., 2013, Lisman and Jensen, 2013, Hyafil et al., 2015, Bonnefond et al., 2017]. Consistent with this notion, PAC has also been reported across distinct brain regions [Sweeney-Reed et al., 2014]. In sum, PAC has been proposed as a canonical mechanism for neural syntax [Buzs´aki, 2010].

Given the growing interest in CFC, and in PAC more specifically, developing adequate and unbiased tools to quantify the posited signatures of neural computations has motivated a number of contributions [Canolty et al., 2006, Penny et al., 2008, Tort et al., 2010]. While these works have met some success, they have also pointed out numerous methodological challenges [Aru et al., 2015, van Wijk et al., 2015] illustrated and discussed in several simulation studies [Tort et al., 2010, Ӧzkurt and Schnitzler, 2011]. Some important limiting factors and potential pitfalls of current used techniques are related to the impact of incorrectly choosing the bandwidth of bandpass filters [Dvorak and Fenton, 2014, Aru et al., 2015], the consequences of applying Hilbert transform on wide-band signals [Chavez et al., 2006, Dvorak and Fenton, 2014], and the potential misidentification of CFC [Hyafil, 2015, Kramer et al., 2008]. The direction we advocate to address these issues is to use a signal model that would fit the data in the sense of the mean squared error, and from which a measure of PAC or CFC could be derived. What all existing metrics have in common is that they do not make use of a signal model, and therefore setting the parameters of the method, such as filtering parameters, can only be driven by how much they lead to a strong PAC/CFC metric. As a consequence, even though current metrics give reasonable estimates of PAC, a legitimate and controlled comparison of methods and parameters, and therefore of the results, is impossible. Additionally, while simulations provide better control, they do not fully solve this issue faced by experimentalists since a simulation may approximate at best, or miss at worst, the real structure of neurophysiological signals. Hence, with the present contribution, we hope to propose a major improvement in CFC/PAC estimation by adopting a principled modeling approach.

To initiate this statistical model-based approach, we propose to consider a signal model which is rich enough to capture time-varying phenomena such as PAC. A model-based approach allows computing the likelihood of a recorded neural signal, which can be interpreted as a measure of the goodness of fit of the model. In the present case, the goodness of fit corresponds to the classical measure of explained variance. Such an evaluation metric is a natural criterion to compare models, and a first step towards an automatic model parameters selection on brain data. Additionally, with a model-based approach, it becomes possible to answer a large variety of practical questions: From a statistical standpoint, one can define which parametrization or which hyper-parameter should be selected, while from a neuroscientific standpoint, one could ask, for instance, whether the instantaneous amplitude of slow oscillations contributes to PAC, or whether the amplitude modulation affects the entire neural spectrum identically. Importantly, our statistical signal modeling approach is not a biophysical one and is thus distinct from previous work [Hyafil et al., 2015, Chehelcheraghi et al., 2017]. Our goal here is to better explain and describe the empirical data themselves in the absence of any assumption regarding the neural mechanisms that have generated them.

To capture PAC in univariate time-series, we propose to use auto-regressive (AR) models, which are stochastic signal models as opposed to deterministic. A canonical deterministic signal model consists in modeling a time-series as a pure sinusoid corrupted by some additive noise. As recently discussed, considering neural oscillations as pure sinusoidal brain responses is an oversimplification of the complex neurophysiological reality [Cole and Voytek, 2017]. AR models do not make the strong assumption of sinusoidality and are rather based on the ability of the model to forecast the signal values. AR models have been successfully used to address multiple problems in neurophysiology such as spectral estimation [Spyers-Ashby et al., 1998], temporal whitening [Mahan et al., 2015], and connectivity measures like Granger causality [Granger, 1988, Valdés-Sosa et al., 2005, Haufe et al., 2010] or coherence [Baccal´a and Sameshima, 2001].

In the present work, we show how driven auto-regressive (DAR) models [Grenier, 2013, Dupré la Tour et al., 2017] can be extended in order to study PAC. Standard AR models are statistically efficient given their low number of parameters, but they are linear, and therefore they cannot directly model non-linear phenomena like PAC. In order to extend AR models to cope with such situations of non-linearity and non-stationarity in signals, various advanced AR models have been proposed in other research fields such as audio signal processing and econometrics [Tong and Lim, 1980, Chan and Tong, 1986, Grenier and Omnes-Chevalier, 1988] (see related non-linear AR models section below). Here, we consider the slow oscillation as the exogenous driver so as to allow the coefficients of the AR model to be time varying, thereby making the instantaneous power spectral density (PSD) of the signal be a function of a slow exogenous time series. By doing so, DAR models are statistical signal models (and not biophysical ones) that do not model PAC explicitly but which are able to capture it.

In what follows, we first present how PAC is commonly analyzed in neural time series while pointing out limitations. We then review the current literature on non-linear AR models beyond neuroscience. We then detail the methodological aspects of DAR models first generically and then specifically for brain data. We describe how we can derive from a DAR model a power spectral density parametrized by an external signal and how it allows to compute so-called comodulograms and estimate directionality of coupling; this enables a fine-grained analysis of spectral modulations observed in PAC. Finally, results show how the comparison of goodness of fit between different models allow to conclude on optimal parameters (e.g. filtering parameters), leading to new neuroscientific insights, such as the wideband or the asymetric spectral properties of the driver. We also report that taking into account the instantaneous amplitude of the driver improves the goodness of fit, which suggests that this information is relevant to the coupling phenomenon and should not be discarded as in most PAC metrics. Last but not least, directionality estimation results reveal some delays between high frequency bursts and the driving low frequency oscillation.

### Notations

We note *y* the signal containing the high-frequency activity, and *x* the signal with slow frequency oscillations, also called the exogenous driver. When a signal *x* results from a band-pass filtering step, we note the central frequency of the filter *f*_*x*_ and the bandwidth ∆*f*_*x*_. The value of the signal *x* at time *t* is denoted *x*(*t*).

### PAC quantification methods and current limitations

To estimate PAC, the typical pipeline reported in the literature consists in four main processing steps:

1. Bandpass filtering is performed in order to extract the narrow-band neural oscillations at both low frequency *f*_*x*_ and high frequency *f*_*y*_;
2. A Hilbert transform is applied to get the complex-valued analytic signals of *x* and *y*;
3. The phase ***ϕ***_*x*_ of the low frequency oscillation and the amplitude *a*_*y*_ of the high frequency signals are extracted from the complex-valued signals;
4. A dedicated approach is used to quantify the correlation between the phase ***ϕ***_*x*_ and the amplitude *a*_*y*_ signals.

The Modulation Index (MI) described in the pioneering work of [Canolty et al., 2006] is the mean over time of the composite signal *z* = *a*_*y*_*e*^****ϕ****_*x*_^. The stronger the coupling between ***ϕ***_*x*_ and *a*_*y*_, the more the MI deviates from zero. This index has been further improved by Ozkurt *et al*. with a simple normalization [Ӧzkurt and Schnitzler, 2011]. Another approach [Lakatos et al., 2005, Tort et al., 2010] has been to partition [0, 2π] into smaller intervals to get the time points *t* when ***ϕ***_*x*_(*t*) is within each interval, and to compute the mean of *a*_*y*_(*t*) on these time points. PAC was then quantified by looking at how much the distribution of *a*_*y*_ differs from uniformity with respect to ***ϕ***_*x*_. For instance, a simple height ratio [Lakatos et al., 2005], or a Kullback-Leibler divergence as proposed by Tort *et al*. [Tort et al., 2010], can be computed between the estimated distribution and the uniform distribution. Alternatively, it was proposed in [Bruns and Eckhorn, 2004] to use direct correlation between *x* and *a*_*y*_. As this method yielded artificially weaker coupling values when the maximum amplitude *a*_*y*_ was not exactly on the peaks or troughs of *x*, this method was later extended to generalized linear models (GLM) using both cos(***ϕ***_*x*_) and sin(***ϕ***_*x*_) by Penny *et al*. [Penny et al., 2008]. Other approaches employed a measure of coherence [Colgin et al., 2009] or the phase-locking value [Lachaux et al., 1999]. All these last three approaches offer metrics which are independent of the phase at which the maximum amplitude occurs. The methods of Tort *et al*. [Tort et al., 2010], Ӧzkurt *et al*. [Ӧzkurt and Schnitzler, 2011], and Penny *et al*. [Penny et al., 2008] will be considered for comparison in our experiments.

As one can see, there is a long list of methods to quantify CFC in neural time series. Yet, a number of limitations which can significantly affect the outcomes and interpretations of neuroscientific findings exist with these approaches. For example, in typical PAC analysis, a systematic bias rises where one constructs the so-called *comodulogram*. A comodulogram is obtained by evaluating the chosen metric over a grid of frequency *f*_*x*_ and *f*_*y*_. This bias emerges from the choice of the bandpass filter, which involves the critical choice of the bandwidth ∆*f*_*y*_. It has been reported several times that to observe any amplitude modulation, the bandwidth of the fast oscillation ∆*f*_*y*_ has to be at least twice as high as the frequency of the slow oscillations *f*_*x*_: *∆f*_*y*_ > 2*f*_*x*_ [Berman et al., 2012, Dvorak and Fenton, 2014]. As a comodulogram uses different values for *f*_*y*_, many studies have used a variable bandwidth, by taking a fixed number of cycles in the filters. The bandwidth is thus proportional to the center frequency: ∆*f_y_ ∝ f*_*y*_. This choice leads to a systematic bias, as it hides any possible coupling below the diagonal *f*_*y*_ = 2*f_x_/α*, where α = ∆*f_y_/f*_*y*_ is the proportionality factor. Other studies have used a constant bandwidth ∆*f*_*y*_; yet this also biases the results towards the low driver frequency *f*_*x*_, considering that it hides any coupling with *f_x_ > ∆f_y_/*2. A proper way to build a comodulogram would be to take a variable bandwidth ∆*f_y_ ∝ f*_*x*_, with ∆*f_y_ >* 2*f*_*x*_. However, this is not common practice as it is computationally very demanding, because it implies to bandpass filter *y* again for each value of *f*_*x*_.

Another common issue arises with the use of the Hilbert transform to estimate the amplitude and the phase of real-valued signals. Such estimations rely on the hypothesis that the signals *x* and *y* are narrow-band, i.e. almost sinusoidal. However, numerous studies have used this technique on very wide-band signals such as the entire gamma band (80-150 Hz) [Canolty et al., 2006] (see other examples in [Chavez et al., 2006]). The narrow-band assumption is debatable for high frequency activity and, consequently, using the Hilbert transform may yield non-meaningful amplitude estimations, and potentially poor estimations of PAC [Chavez et al., 2006, Dvorak and Fenton, 2014]. Note also that, in this context, wavelet-based filtering is equivalent to the Hilbert transform [Quiroga et al., 2002, Bruns, 2004], and therefore does not provide a more valid alternative option.

Besides these issues of filtering and inappropriate use of Hilbert transforms, Hyafil [Hyafil, 2015] also warned that certain choices of bandwidth ∆*f*_*y*_ might mistake phase-frequency coupling for PAC, or create spurious amplitude-amplitude coupling; see also the more recent work in [Aru et al., 2015] for discussion and more practical recommendations for PAC analysis.

Here we advocate that the DAR models detailed in the next sections address a number of the limitations just mentioned. They do *not* use bandpass filter or Hilbert transform on the high frequencies *y*. They introduce a measure of *goodness of fit*, through the use of a probabilistic signal model whose quality can be assessed by evaluating the *likelihood* of the data under the model. In practice, the likelihood quantifies how much variance of the signal can be explained by the model, and is similar to the *R*^2^ coefficient in generalized linear models (GLM). To the best of our knowledge, the only related model-based approach to measure PAC used GLM [Penny et al., 2008]. With GLM, however, the modeling part is done independently on each signal *y*_*f*_, which is the band-pass filtered version of *y* around frequency *f*. For each of these frequencies *f* a different model is fitted. By doing so, a GLM approach cannot model the wide-band signal *y* as it is limited to multiple estimations frequency by frequency bin. This largely limits the use of the likelihood to compare models or parameters. On the contrary, we propose to model *y* globally, without filtering it in different frequency bands.

To conclude this section, and to position this work in the broader context of modeling approaches for neuroscience data, we would like to stress that our proposed method can be considered as an encoding model for CFC, as opposed to a decoding model [Kay et al., 2008, Naselaris et al., 2011, Huth et al., 2016]. Indeed, our model reports how much empirical data can be explained and by doing so enables us to test neuroscience hypotheses in a principled manner [Naselaris et al., 2011].

### Non-linear AR models beyond Neuroscience

The literature on the use of non-linear auto-regressive (AR) models is quite large and covers fields such as audio signal processing and econometrics. For instance, AR models with conditional heteroskedasticity (ARCH [Engle, 1982], GARCH [Bollerslev, 1986]) are extremely popular in econometrics where they are used to model signals whose overall amplitude varies as a function of time. Here, however, in the context of CFC and PAC, one would like to model variations in the spectrum itself, such as shifts in peak frequencies (a.k.a. frequency modulations) or changes in amplitude only within certain frequency bands (a.k.a. amplitude modulations). To achieve this, one idea is to define a linear AR model, whose coefficients are a function of time and change slowly depending on a *non-linear* function of the signal.

The first models based on this idea are SETAR models [Tong and Lim, 1980], which switch between several AR models depending on the amplitude of the signal with respect to some thresholds. To get a smoother transition between regimes, SETAR models have inspired other models like EXPAR [Haggan and Ozaki, 1981] or STAR [Chan and Tong, 1986], in which the AR coefficients change continuously depending on a non-linear function of the past of the signal.

These models share the same underlying motivation as the DAR models described below but, crucially, DAR models can be designed and parametrized to capture PAC phenomena independently of the phase in the driving signal at which the high frequency content is the strongest. In other words, DAR models can work equivalently well if the high frequency peaks are in the troughs, the rising phase, the decreasing phase or the peaks of the low frequency driving signal. Moreover, as DAR models do not require to infer the driving behavior from the signal itself and rather rely on the prior knowledge of the slow oscillation, the inference is significantly faster and more robust.

## Methods

In this section, we first define our statistical signal model, explain how we estimate its parameters on a signal *y* and its driver *x*, and demonstrate how one can infer hyper-parameters by comparing the likelihood of several models. Then, we detail how to use this model on neurophysiological signals by presenting the preprocessing steps and showing how to make the model invariant to the phase of the coupling. We also detail how to express the power spectral density (PSD) conditionally to the driver’s phase, which allows for a fine-grained analysis of the signal’s spectral properties and to build comodulograms. Finally, we present our protocol to simulate signal with CFC and the empirical datasets used to validate our method.

### DAR models - General presentation

#### Model definition

Let *y* be a univariate locally stationary signal, as defined in [Dahlhaus, 1996]. An auto-regressive (AR) model specifies that *y* depends linearly on its own *p* past values, where *p* is the *order* of the model:

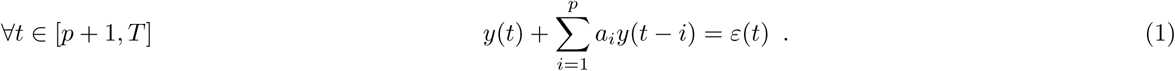

Here, *T* is the length of the signal and ε is the *innovation* (or *residual*) modeled with a Gaussian white noise: 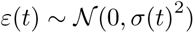. To extend this AR model to a non-linear model, one can assume that the AR coefficients *a*_*i*_ are non-linear functions of a given exogenous signal *x*, here called the *driver*:

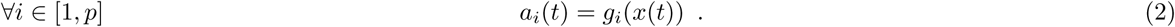

What was proposed in [Grenier, 2013, Grenier, 1983] is to consider that the non-linear functions *g*_*i*_ are polynomials:

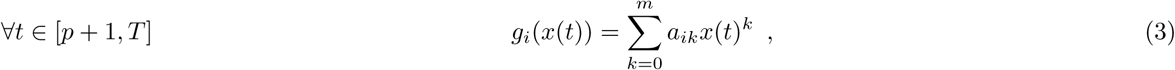

where *x(t)*^*k*^ is *x(t)* to the power *k*. Note that one could imagine more complex models for the *g*_*i*_, yet empirical evidence confirms that polynomials are rich enough for the different neurophysiological signals considered in this work (see S1 Fig).

Inserting the time-dependent AR coefficients (2) into the AR model (1), we obtain the following equation:

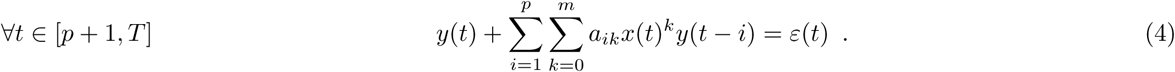

This can be simplified and rewritten as:

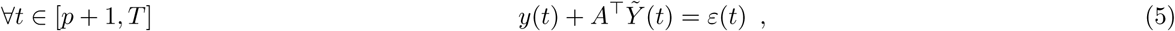

where 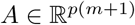 is a one-dimensional vector composed of the scalars 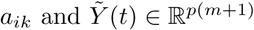 is a one-dimensional vector composed of the regressors *x*(*t*)^*k*^*y*(*t — i*). We note 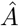 the estimated value of *A* and *A*^*T*^ its transpose.

We also consider a time-varying innovation variance σ(*t*)^2^ driven by *x*. This corresponds to the assumption that the power of the signal at time *t* depends on the driver at this same instant. Since the standard deviation is necessarily positive, we use the following polynomial model for its logarithm:

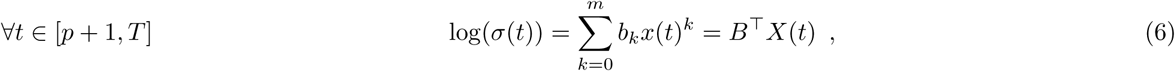

where 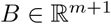 is a one-dimensional vector composed of the scalars *b*_*k*_.

This model is called a driven auto-regressive (DAR) model [Dupré la Tour et al., 2017]. DAR models are thus a natural extension of well-known AR models and enable analyzing non-stationary or non-linear spectral phenomena observed in neurophysiological time-series. One methodological concern is that the AR coefficients cannot be arbitrary as they should define stable models (see [Proakis and Manolakis, 1996], section 3.6). In our case there is an extra difficulty because they are changing over time. To guarantee that each set of coefficients defines a stable filter, it is possible to re-parametrize the model. This requires significantly more complex derivations and leads to a slightly slower optimization, while providing equivalent empirical observations. For simplicity, this manuscript will focus on the simplest model dedicated to CFC analysis, and we refer to [Dupré la Tour et al., 2017, Grenier and Omnes-Chevalier, 1988] for readers interested in more details.

#### Model estimation

DAR model equation (4) is non-linear with respect to the given signals *x* and *y*, yet it is linear with respect to the regressors *x*(*t*)^*k*^*y*(*t — i*). Therefore after computing the regressors, it is possible to obtain an analytical expression of the parameters in *A*.

As the innovation ε(*t*) is assumed to be a Gaussian white noise, the model likelihood *L* is obtained via:

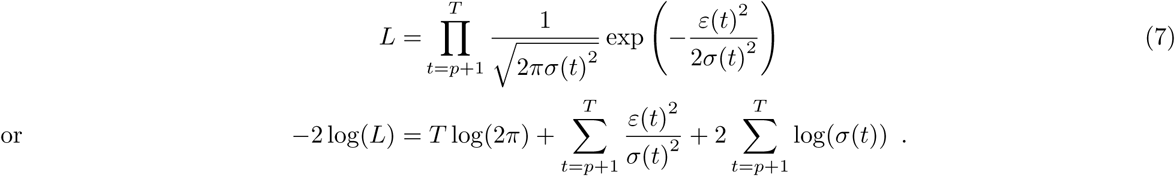

DAR models are estimated by likelihood maximization. Here, if the innovation variance σ(*t*)^2^ is considered fixed, maximizing this function boils down to minimizing the sum of squares of the ε(*t*). Since ε(*t*) is the residual, the inference of the parameters amounts to maximizing the variance explained by the model. To start the maximization, we first assume that the innovation variance is fixed and equal to the signal’s empirical variance. The likelihood then reads:

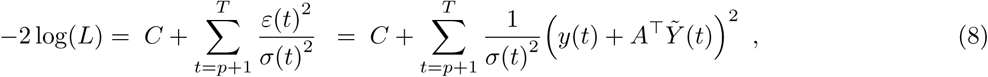

where *C* is a constant. We can thus estimate the DAR coefficients 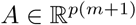 by solving the following linear system (a.k.a. normal equations):

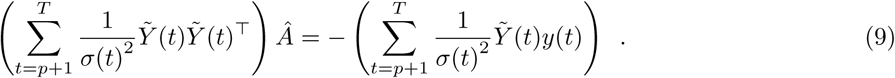

Given 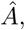 one can then estimate the vector 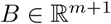 from the residual ε(*t*), maximizing the likelihood using off-the-shelf optimization techniques adapted to differentiable problems. We use a Newton-Raphson procedure (see more details in [Grenier, 2013]). The instantaneous variance is then updated with *B* using (6). One can then iterate between the estimation of the coefficients *A* and *B*. In our experiments, we observed that one or two iterations are sufficient.

#### Model selection

DAR models are probabilistic, which offers a significant advantage over traditional PAC metrics. Indeed, one can evaluate the likelihood of the model *L* (7), which quantifies the *goodness of fit* of the data given the model, and select the hyper-parameters that maximize it. To compare models with different number of hyper-parameters (i.e. degrees of freedom), different information criteria can be used such as the Akaike Information Criterion (AIC) [Akaike, 1998] or the Bayesian Information Criterion (BIC) [Schwarz et al., 1978], which read respectively AIC = −2 log(*L*) + 2*d* and BIC = −2 log(*L*) + *d* log(*T*). The integer *d* is the number of degrees of freedom of the model, which for DAR models equals to *d* = (*p* + 1)(*m* + 1). The BIC has been used extensively for hyper-parameter selection in AR models, and enables the joint estimation of both *p* and *m* (see S2 Fig for simulation results).

As the statistical inference with a DAR model is very fast, one can easily estimate models on a full grid of hyper-parameters (*p* and *m*) and keep the model that leads to the lowest BIC. Another alternative used in our experiments is the use of cross-validation (CV), consisting of splitting the recorded signals into two sets: a training signal to estimate parameters, and a test signal to evaluate the likelihood of new data. In practice, CV requires that a large amount of data be left for model testing, and that the two datasets be separated by a minimum delay to ensure data independence [Arlot et al., 2010]. In S3 Fig, we present model selection based on cross-validation, applied on real signals.

#### Model’s power spectral density

AR models are mainly used for spectral estimation. Since an AR model is a linear filter that is fitted to somehow whiten a particular signal *y*, the power spectral density (PSD) of this filter is close to the inverse of the PSD of the signal, providing a robust spectral estimation of the signal. For a linear AR model, the PSD at a frequency *f* is given by:

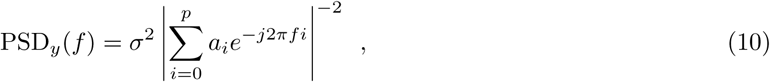

where *j*^2^ = –1 and *a*_0_ = 1. This estimation is perfect when *p* → ∞, but in practice, using *p ϵ* [10, 100] gives satisfying results in most applications.

AR models have been successfully used for spectral estimation in a wide variety of fields, such as geophysics, radar, radio astronomy, oceanography, speech processing, and many more [Kay and Marple, 1981]. Importantly, they can be applied on any signal, even deterministic, as they can be seen as linear predictions based on the *p* past elements, independently from the way the signal was generated. However, in the specific case of a noisy sum of perfect sinusoids, the peak amplitudes in AR spectral estimates are proportional to the square of the power [Kay and Marple, 1981], unlike conventional Fourier spectral estimates where the peak amplitudes are proportional to the power. Pisarenko’s method [Pisarenko, 1973], later extended in the MUSIC algorithm [Schmidt, 1986], might be more suited for spectral estimation of such ‘ideal’ signals.

However, as neurophysiological signals are not perfect sinusoids, AR model are well suited for their analysis. For a good overview of spectral estimation techniques, including AR models, we refer the reader to Kay and Marple [Kay and Marple, 1981].

Interestingly, AR models yield a spectral estimation with an excellent frequency resolution. Here, frequency resolution is not defined as the distance between two points in the PSD, which would go to zero as we increase zero-padding in the AR filter. Rather, the frequency resolution is defined as the smallest distance that can be detected between two spectral peaks. With this definition, spectral estimation with AR models has a better frequency resolution than Fourier-based methods [Marple, 1977].

Another benefit of spectral estimation with AR models is the theoretical guarantee in term of maximum entropy. Maximizing the entropy of the spectrum consists in introducing as little as possible artificial information in the spectrum. Assuming that we have *p* auto-correlations of a wide-sense stationary signal, maximizing the entropy leads to the maximum likelihood estimate of an AR model [Ables, 1974]. In other words, the maximum likelihood estimate of an AR model is the best spectral estimator in term of maximal entropy.

In the case of DAR models, the PSD is a function of the driver’s value *x*, so we note it PSD_*y*_(*x*). For a given complex driver’s value *x*_0_, we compute the AR coefficients *a*_*i*_(*x*_0_) using Eq. (2), along with the innovation’s standard deviation σ(*x*_0_) using Eq. (6). Since AR models with time-varying coefficients are locally stationary [Dahlhaus, 1996], we can compute the PSD at a frequency *f* with:

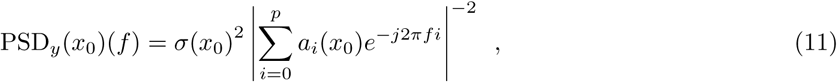

where *j*^2^ = –1 and *a*_0_(*x*_0_) = 1. We thus have a different PSD for each driver value *x*_0_, i.e. for each time instant, as the driver *x*(*t*) fluctuates in time. Note that the PSD is a function of the driver value and not only its phase, unless the driver has been normalized in amplitude. The varying AR coefficients *a*_*i*_(*t*) parametrize the fluctuations of the shape of the PSD, whereas the varying innovation’s standard deviation σ(*t*) is responsible for the absolute fluctuations in power over all frequencies.

### DAR models for CFC estimations on brain data

#### Data preprocessing: Extracting the driver

In this section, we describe the preprocessing applied to the raw signal *z* to obtain the signal *y* and the driver *x*.

First, the raw signal was downsampled to an appropriate sampling frequency *f*_*s*_ (we used 333 Hz and 625 Hz in our examples), as we only consider frequencies up to *f*_*s*_/2. Power line noise and its harmonics were removed. As power line noise fluctuates over time, the frequency of the power line noise was estimated by projecting the signal onto sinusoids, and the maximum-power projection was subtracted from the raw signal. When necessary, a high-pass filter was used to detrend the signal (typically at 1 Hz) [Bigdely-Shamlo et al., 2015].

A bandpass filter was then used to extract the driver *x*. The center frequency *f*_*x*_ and bandwidth ∆*f*_*x*_ for this filter will be discussed in details in the Results section. The filter equation is:

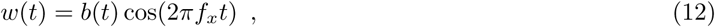

where *b*(*t*) is a Blackman window of order ⌊1.65 ∗ *f*_*s*_/∆*f*_*x*_⌋ ∗ 2 + 1, chosen to have a bandwidth of ∆*f*_*x*_ at −3 dB (the filter attenuation is 50% at *f*_*x*_ ± ∆*f*_*x*_/2). The filter is zero-phase since it is symmetric.

Then, the driver *x* is subtracted from the raw signal *z* to create the signal *y* = *z − x*. Note that the signal *y* now contains a frequency gap around *f*_*x*_, which can be a nuisance for the AR estimate that provides a compact model for the broad band power spectrum density of the signal. To solve this issue, we fill this gap by adding a Gaussian white noise filtered with the filter *w*, and adjusted in energy to have a smooth power spectral density (PSD) in *y*. Such a preprocessing step is also commonly used when working with multivariate auto-regressive models (MVAR) on neurophysiological signals corrected for power line noise.

Finally, we whiten *y* with a linear AR model, by applying the inverse AR filter to the signal. This temporal whitening step is not necessary, yet it reduces the need for high order *p* in DAR and therefore reduces both the computational cost and the variance of the model. After this whitening step, the PSD of *y* is mostly flat, and DAR models only contain the modulation of the PSD. Fig. 1 shows a time sample of a preprocessed signal, taken from a rodent striatal LFP signal (see below for a description of the data). One can see how the slowing varying driving signal follows the original raw signal and how the high frequencies bursting on the peak of the slow oscillation remain in the processed signal *y*. The general processing pipeline is summarized in Fig. 2(a).

**Fig 1.**
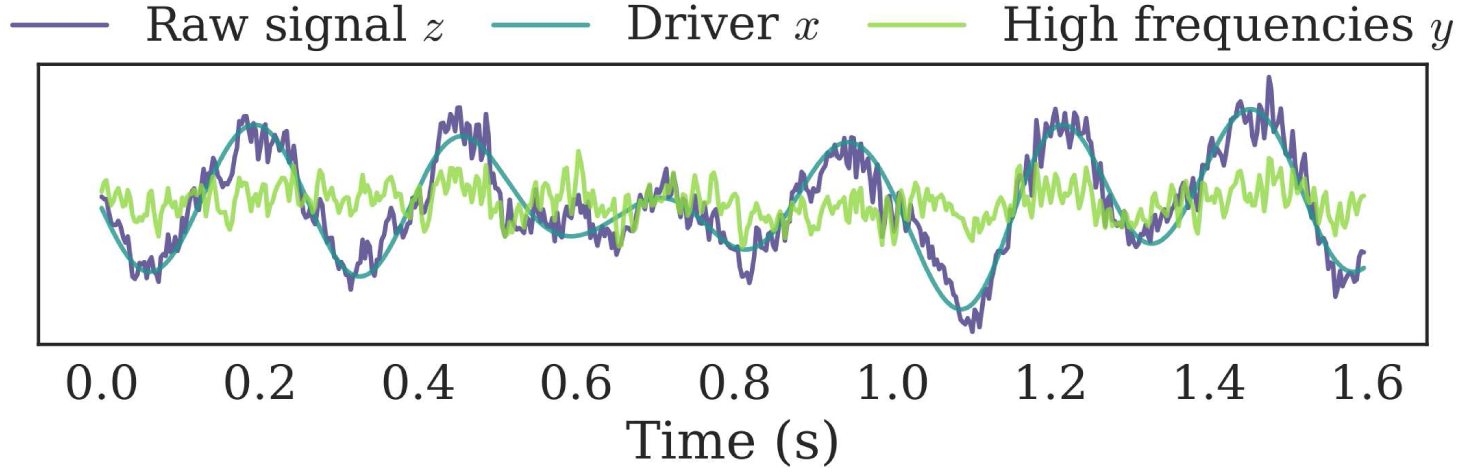
Example of a preprocessed signal. The driver *x* is extracted from the raw signal *z* with a bandpass filter, and then subtracted from *z* to give *y*. A PAC effect is present as we see stronger high frequency oscillations at the peaks of the driver. The signal *y* is presented here as it looks before the temporal whitening step.

**Fig 2.**
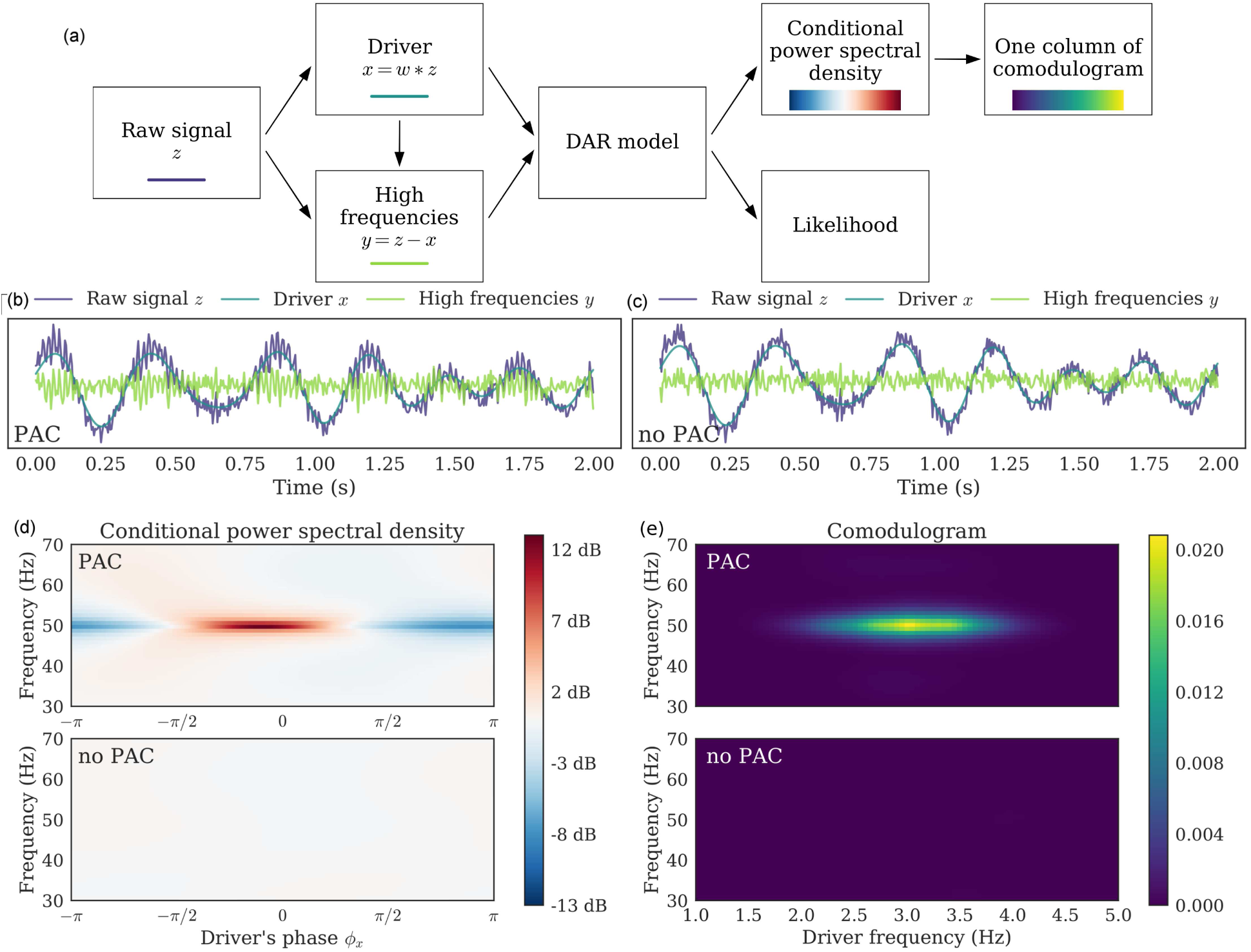
Example on simulated signals: Pipeline, signal, conditional PSD, and comodulogram. (a) Pipeline of the method. We applied it with (*p, m*) = (10, 1) on two simulated signals: (b) Simulated signal with PAC and (c) Simulated signal without PAC. (d) From a fitted model, we computed the PSD conditionally to the driver’s phase ***ϕ***_*x*_; each line is centered independently to show amplitude modulation. PAC can be identified in the fluctuation of the PSD as the driver’s phase varies: around 50 Hz, the PSD has more power for one phase than for another. This figure corresponds to a single driver’s frequency *f*_*x*_ = 3.0 Hz. (e) Applying this method to a range of driver’s frequency, we build a comodulogram, which quantifies the PAC between each pair of frequencies.

In the case of analyzing CFC between two different channels *A* and *B*, we can simply perform the previous steps on both raw signals *z_A_* and *z_B_*, and fit a DAR model on *y_A_* driven by *x*_*B*_. It could also be possible to not subtract *x*_*A*_ from *z*_*A*_, but if the low frequency bands *x*_*A*_ and *x*_*B*_ are assumed to be uncorrelated, which might not be the case. In such case, the other preprocessing steps should still be performed on *z*_*A*_.

#### Model selection: Optimizing the filtering parameters

During the preprocessing, the driver *x* is extracted from the preprocessed signal *z* through a band-pass filter, with center frequency *f*_*x*_ and bandwidth ∆*f*_*x*_. To select automatically these two parameters, we perform a grid-search and select the driver which yields the highest likelihood. Importantly, the highest likelihood does not necessarily correspond to the highest PAC score, but rather to the model with the highest goodness of fit or explained variance. This detail is crucial, as optimizing for the best fit is a legitimate data-driven approach, whereas optimizing for the highest PAC score is statistically more questionable. One additional benefit of our approach is that other types of filters can be compared using a common methodological approach.

For a fair comparison of the drivers, we need to evaluate the model fitting on the exact same signal *y*. To do so, we remove from *y* all the possible drivers tested on this grid-search using a high-pass filter above maximum center frequency *f*_*x*_. Once again, we fill the frequency gap by adding a Gaussian white noise filtered with a low-pass filter complementary to the high-pass filter.

#### Phase invariance: Removing the bias using a complex driver

We consider PAC where high frequencies are coupled with a low frequency driving signal. We denote by ***ϕ***_0_ the preferred phase of the coupling, i.e. the phase of the driver that corresponds to the maximum amplitude of the high frequencies. When ***ϕ***_0_ = 0, the high frequency bursts happen in the peaks of the driver, whereas when ***ϕ***_0_ = π, they happen in the troughs.

The driver extracted as described in previous subsection is real-valued, and the filter used to extract it is based on a cosine. As the cosine has the same value for ****ϕ**** and π *− ****ϕ*****, the PAC estimation is biased and underestimated when ****ϕ****_0_ is not equal to 0 or *π*. Indeed, the model mixes the contribution of ****ϕ**** and π *****ϕ*****, which attenuates the effect. This bias was already reported in PAC estimation by [Bruns and Eckhorn, 2004], and the solution proposed by [Penny et al., 2008] was to use both cosine and sine to have PAC estimates that are invariant to the preferred phase ****ϕ****_0_ of the low frequency oscillation.

The same technique can be used on the DAR models. We not only filter the raw signal with *w*_1_(*t*) = *b*(*t*) cos(2π*f_x_t*) to obtain *x*_1_, but also with *w*_2_(*t*) = *b*(*t*) sin(2π*f_x_t*) to obtain *x*_2_, creating a complex-valued driver *x* = *x*_1_ + *jx*_2_ where *j* denotes the complex number, *j*^2^ = –1. With a complex-valued driver, DAR models are naturally extended by adding more regressors:

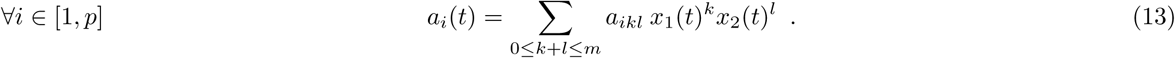

Instead of *p*(*m* + 1) regressors, we now have *p*(*m* + 1)(*m* + 2)*/*2 regressors, and the number of degrees of freedom of the model is now *d* = (*p* + 1)(*m* + 1)(*m* + 2)*/*2. Typical values for *m* are below 3, so the number of parameters stays within a reasonable range despite the squared dependence in *m*. The number of parameters in the model remains much lower than the number of time samples and the estimation problem stays therefore well-posed. The innovation variance model is also updated accordingly:

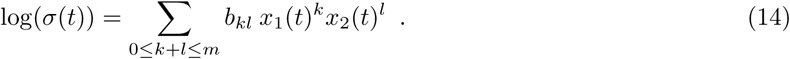

A validation result is presented in the supporting information (S4 Fig), showing that indeed it removes the preferred-phase bias.

#### Conditional spectrum: Deriving the PSD from the model

After preprocessing and model estimation, we can compute the conditional power spectral density (PSD) of the DAR model, as described earlier. In Fig. 2(b and c), one can see two simulated signals: one with PAC on the left (b) and one without PAC on the right (c). The simulation process, which does not make use of DAR model, will be detailed and discussed in a following section. In Fig. 2(d), one can see an example of PSD_*y*_(*x*_0_), with artificial driver’s values *x*_0_ = ρ exp(*j****ϕ*****). In practice we set ρ to the median of the empirical driver’s amplitude, ρ = median(|*x*|), and *****ϕ***** spans the entire interval [–*π, π*]. Computing the PSD on a circle allows to visualize the PSD fluctuations with respect to the driver’s phase, giving a representation similar to other PAC metrics. The PAC is identified by the modulation of the spectrum with respect to the driver’s phase ****ϕ****. Other trajectories could be chosen to visualize also the dependency on the driver’s amplitude. Note that with DAR models, the amplitude modulation of *y* is modeled jointly on the entire spectrum, as opposed to frequency band by frequency band in most other PAC metrics.

Using the fast Fourier transform (FFT) in Eq. (11), we are able to compute the PSD for a broad range of frequencies *f* ϵ [0, *f*_*s*_/2]. As model estimation is also very fast, we obtain the conditional PSD very quickly: With (*p, m*) = (10, 1) on a signal with *T* = 10^5^ time points, the entire DAR estimation and PSD computation lasts 0.12 s. With larger values (*p, m*) = (90, 2) as selected by cross-validation on the rodent striatal dataset, it lasts 1.34 s.

#### Comodulogram: Quantifying the modulation at each frequency

We now detail how comodulograms can be derived from DAR models. To quantify the coupling at a given frequency *f*, we compute PSD_*y*_(*x*_*c*_)(*f*) for a range of *n* artificial driver’s values *x*(*k*) located on a circle of radius ρ = median(|*x*(*t*)|), and we normalize it to sum to one:

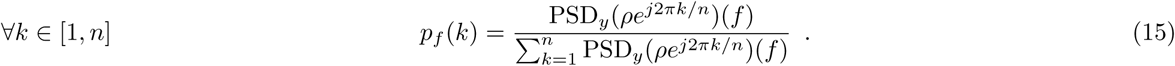

Then, we use the same method as [Tort et al., 2010] to measure the fluctuation of *p*_*f*_ (*k*): we compute the Kullback-Leibler divergence between *p*_*f*_ (*k*) and the uniform distribution *q*(*k*) = 1*/n*, and we normalize it with the maximum entropy log(*n*):

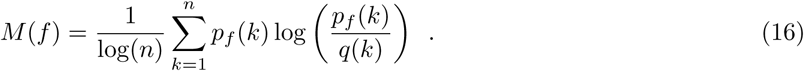

This metric *M* is between 0 and 1 and allows us to quantify the non-flatness of *p*_*f*_ (*k*), i.e. the amplitude modulation, frequency by frequency. We can then build a *comodulogram*, a figure commonly used in the literature which depicts the PAC metric for a grid of frequency (*f_x_, f*). When derived from DAR models as described here, the comodulogram is not affected by the systematic biases presented in the introduction.

Fig. 2(e) shows the comodulogram of the two simulated signals, one with and one without PAC. As expected the DAR model reports no coupling when there is none, and reports here a strong coupling between 3 and 50 Hz when it is actually present.

#### Directionality: Estimating a delay between the signal and the driver

The DAR model in Eq. (4) uses the driver at time *t* to parametrize the AR coefficients with the assumption that, in PAC, high frequency activity is modulated by the driving signal with no delay. However, one could legitimately ask whether slow fluctuations in oscillatory activity precedes the amplitude modulation of the fast activity or, conversely, whether high frequency activity precedes low frequency fluctuations. This is what we will refer to as *directionality estimation*.

To assess the problem of directionality estimation with DAR models, we capitalize on the likelihood function. After observing some PAC between *y* and *x*, we introduce a delay τ in the driver so as to estimate the goodness of fit of a DAR model between *y* and *x_τ_*, where *x_τ_* (*t*) = *x*(*t − τ*). Using a grid of values for the delay, one can report the value τ_0_ that corresponds to the maximum likelihood. In practice, as we use non-causal filters to extract the driver, and because the AR model uses samples from the past of the *y*, one might wonder if there is any positive or negative bias in our delay estimation. To address this issue, we apply our analysis on both forward and time-reversed signals, and sum the two models log-likelihood. In this way, the filtering bias is strongly attenuated because it similarly affects both forward and time-reversed models. If the best delay is positive, it means that the past driver yields a better fit than the present driver, i.e. that the slow oscillation precedes the amplitude modulation of the fast oscillation. Inversely, a negative delay means that the amplitude modulation of the fast oscillation follows changes in the slow oscillation.

It is worth noting that the estimation of the delay τ_0_ shall not be considered as an alternative way to estimate the preferred phase ****ϕ****_0_ defined in a previous section. Although the preferred phase and the delay would be identical should the driver be a perfect stationary sinusoid, in non-stationary neural systems in which the driver’s instantaneous frequency may fluctuate with time, the preferred phase and delay will likely differ. Such a scenario can occur either because signal waveforms are not perfect sinusoids [Cole et al., 2016] or because the driver actually changes [Tort et al., 2008]. In other words, τ_0_ is a time delay which is identical at all time and corresponds to different phase shifts ****ϕ****(*t*) = τ_0_ ∗ (2π*f* (*t*)) which depend on the instantaneous frequency *f* (*t*). On the contrary, ****ϕ****_0_ is a phase shift that is constant over time and which corresponds to different time delays τ (*t*) = ****ϕ****_0_*/*(2π*f* (*t*)) which also depend on the instantaneous frequency. Fig. 3 illustrates these specific points and disentangles the two distinct notions.

**Fig 3.**
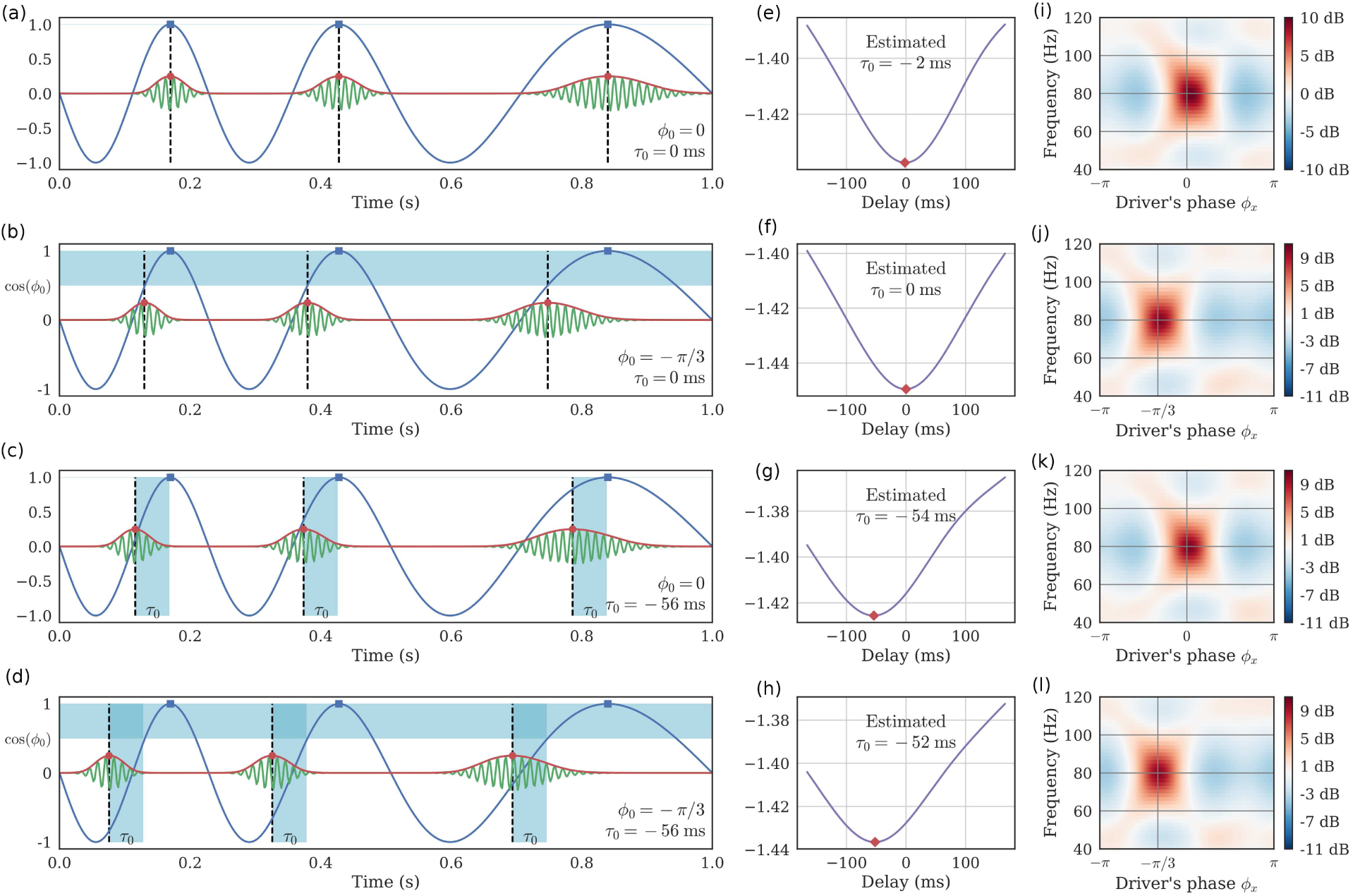
Preferred phase and temporal delay are different. The temporal delay τ is distinct from the preferred phase ****ϕ****_0_. (a) When both are equal to zero, the high frequency bursts happen in the driver’s peaks. (b) When τ = 0 and ****ϕ**** ≠ 0, the bursts are shifted in time with respect to the driver’s peaks, and this shift *varies* depending on the instantaneous frequency of the driver. (c) When τ ≠ 0 and ****ϕ**** = 0, the bursts are shifted in time with respect to the driver’s peaks, and this shift is *constant* over the signal. In this case, note how the driver’s phase corresponding to the bursts varies depending on the instantaneous frequency of the driver. (d) τ and ****ϕ****_0_ can also be both non-zero. (e-h) Negative log-likelihood of DAR models, fitted with different delays between the driver and the high frequencies. The method correctly estimates the delay even when ****ϕ****_0_ 0. (i-l) PSD conditional to the driver’s phase, estimated through a DAR model with the best estimated delay. The maximum amplitude occurs at the phase ****ϕ****_0_.

Like most PAC metrics, DAR models are invariant with respect to the preferred phase. Both the strength of the coupling and the model likelihood are unchanged with respect to ****ϕ****_0_. On the contrary, all PAC metrics including DAR models are strongly affected by a time delay τ when the driver is not a perfect sinusoid. This attenuates the coupling and can potentially modify artificially the preferred phase. When the delay is too large, all metrics would measure zero coupling. This is in fact what justifies the use of surrogate techniques that introduce a large time shift to quantify the variance of the measure in the absence of coupling [Canolty et al., 2006, Tort et al., 2010, Aru et al., 2015]. Fig. 3(e-h) provides the delay estimated with DAR models, obtained by maximizing the likelihood over a grid of delays, as explained previously. In Fig. 3(i-l), we show the conditional PSD of the model obtained with the best estimated delay as well as the estimated preferred phase ****ϕ****_0_. Using the likelihood, our method is thus able to estimate this delay, thus improving PAC estimation and extending PAC analysis.

The question of delay estimation has been previously addressed in the literature with measures borrowed from information theory. For instance, transfer entropy (TE) [Schreiber, 2000, Wibral et al., 2013, Park et al., 2015] has been adapted to PAC by applying it on the driver and the envelop of the fast oscillations [Besserve et al., 2010]. The TE from a signal *a* to a signal *b* is defined as the mutual information between the present of *b* and the past of *a*, conditioned on the past of *b*. It is similar in this way to Granger causality (GC) [Granger, 1969], which also compares how the present of *b* can be better predicted by the past of both *a* and *b*, than by only the past of *b*, using AR models. Note that when variables follow a Gaussian distribution, TE and GC are actually equivalent [Barnett et al., 2009]. In the case of PAC, both TE and GC need to be evaluated on the driver *x* and the *envelope* of a bandpass filtered *y*_*f*_. Interestingly, GC has been shown to fail when the signal-to-noise ratios of the two signals are different [Nolte et al., 2010], which is often the case for the driver and the envelop of the high frequencies. Hence, it remains to be investigated how TE/GC compares with DAR models in terms of performances.

Our method is also different from these other methods, since we model directly *y* and not its envelop, making our method more specific to PAC. This is made possible since DAR models use the delayed driver *x_τ_* not to predict *y*, but to modulate how *y* is predicted from its own past. PAC directly arises from this non-linear interaction in DAR models. Inherently to this PAC specificity, our method is also asymmetrical with respect to *x* and *y*.

Delay estimation in PAC has also been addressed with another method called Cross-Frequency Directionality (CFD) [Jiang et al., 2015], which is based on the phase slope index (PSI) [Nolte et al., 2008]. PSI is a measure of the phase slope in the cross-spectrum of two signals, and is also used to infer causal relations between signals. CFD adapts this measure by applying it on the driver and the envelop of the fast oscillations. Contrary to TE, CFD is not designed to measure a delay; thus, we modified it so as to compare it *qualitatively* to our approach (see Results).

If such delay estimation results may not reflect pure causality [Aru et al., 2015], for instance because of the transitivity of correlation, they nevertheless improve the analysis of PAC going one step further by estimating the delay between the coupled components.

## Data

### Simulated signals

In our experiments, methods comparisons and model evaluations, we make an extensive use of simulated signals showing some PAC. The simulations are generated as follows: we simulate a coupling between *f*_*x*_ = 3 Hz and *f*_*y*_ = 50 Hz, with a sampling frequency *f*_*s*_ = 240 Hz, during *T* time points (*T* depends on the experiment). We do not use a perfect sinusoidal driver, as using such an ideal oscillatory signal is over-simplistic. Drivers can indeed have a larger band as suggested next by our experiments. Finally, strong empirical evidence suggests that neurophysiological signals have complex morphologies that can be overlooked when studied as ideal sinusoids [Cole and Voytek, 2017].

To construct a time-varying driver peaking at 3 Hz, we bandpass filter a Gaussian white noise at a center frequency *f*_*x*_ = 3 Hz and with a bandwidth ∆*f*_*x*_ = 1 Hz, using the same filter detailed in the preprocessing section. The smaller the bandwidth, the closer the driver is to a perfect sinusoid. We normalize the driver to have unit standard deviation σ_*x*_ = 1. We then modulate the amplitude *a*_*y*_ of a sinusoid at, for example, *f*_*y*_ = 50 Hz, using a sigmoid on the driver *x*:

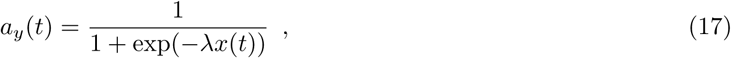

with a sharpness set to λ = 3. By doing so, the amplitude varies between 0 and 1 depending on the driver’s value. We normalize the signal *y* to have a standard deviation σ_*y*_ = 0.4. We then add to the signal *y* both the driver *x* and a Gaussian white noise with a standard deviation σ_ε_ = 1. Note that this simulation procedure is not based on a DAR model. In other words, we do not validate our model using signals that fall perfectly into the category of stochastic signals that are synthesized with a DAR model.

### Empirical datasets

We tested DAR models on three different datasets, with invasive recordings in humans and rodents:

1. A rodent local field potential (LFP) recording in the dorso-medial striatum from [Dallérac et al., 2017](supplementary figure 2), (1800 seconds down-sampled at 333 Hz)
2. A rodent LFP recording in the hippocampus from [Khodagholy et al., 2015] collected during rapid eye movement (REM) sleep (100 seconds down-sampled at 625 Hz)
3. A human electro-corticogram (ECoG) channel in auditory cortex from [Canolty et al., 2006] (730 seconds down-sampled at 333 Hz)

We refer the reader to corresponding articles for more details on the recording modalities of these neurophysiological signals.

## Results

We now illustrate the benefits of DAR models using simulated data and neurophysiological recordings in rats and humans. Our primary goal is to demonstrate that assessing the validity of the model through its likelihood is a statistically principled approach that allows to reveal new insights on empirical data with interesting neuroscientific consequences.

### Model selection and data-driven estimation of the driving signal

In this section, we present the outcome of using the model selection procedure to estimate the best filtering frequency *f*_*x*_ and bandwidth ∆*f*_*x*_ to extract the driver *x*. We first describe the outcome on simulated signals (ground truth) and then on empirical datasets.

#### Simulation study: Validating the approach

To validate our approach, we tested on simulated signals whether both center frequency *f*_*x*_ and bandwidth

*∆f*_*x*_ were correctly estimated. For this, we simulated signals using *f*_*x*_(*simu*) = 4 Hz and

*∆f*_*x*_(*simu*) ∈ [0.2, 0.4, 0.8, 1.6, 3.2] Hz. To mimic the duration of our three real signals, we used *T* = ⌊100*f*_*s*_⌋ time points, for a duration of 100 seconds.

As the noise mainly affected the amplitude modulation but barely altered the driver, we also added a Gaussian white noise low-pass filtered at 20 Hz, and scaled to have a PSD difference of 10 dB at *f*_*x*_ = 4 Hz between the driver and the noise. In this way, the driver was also altered by noise.

Given the simulated signal, we performed a grid-search over *f*_*x*_ and ∆*f*_*x*_, extracting the driver, fitting the model and computing the likelihood of the model. Results are presented in Fig. 4. As expected, the center frequency *f*_*x*_ was correctly estimated for all bandwidths ∆*f*_*x*_. More importantly, the negative log-likelihood was minimal at the correct simulated bandwidth.

**Fig 4.**
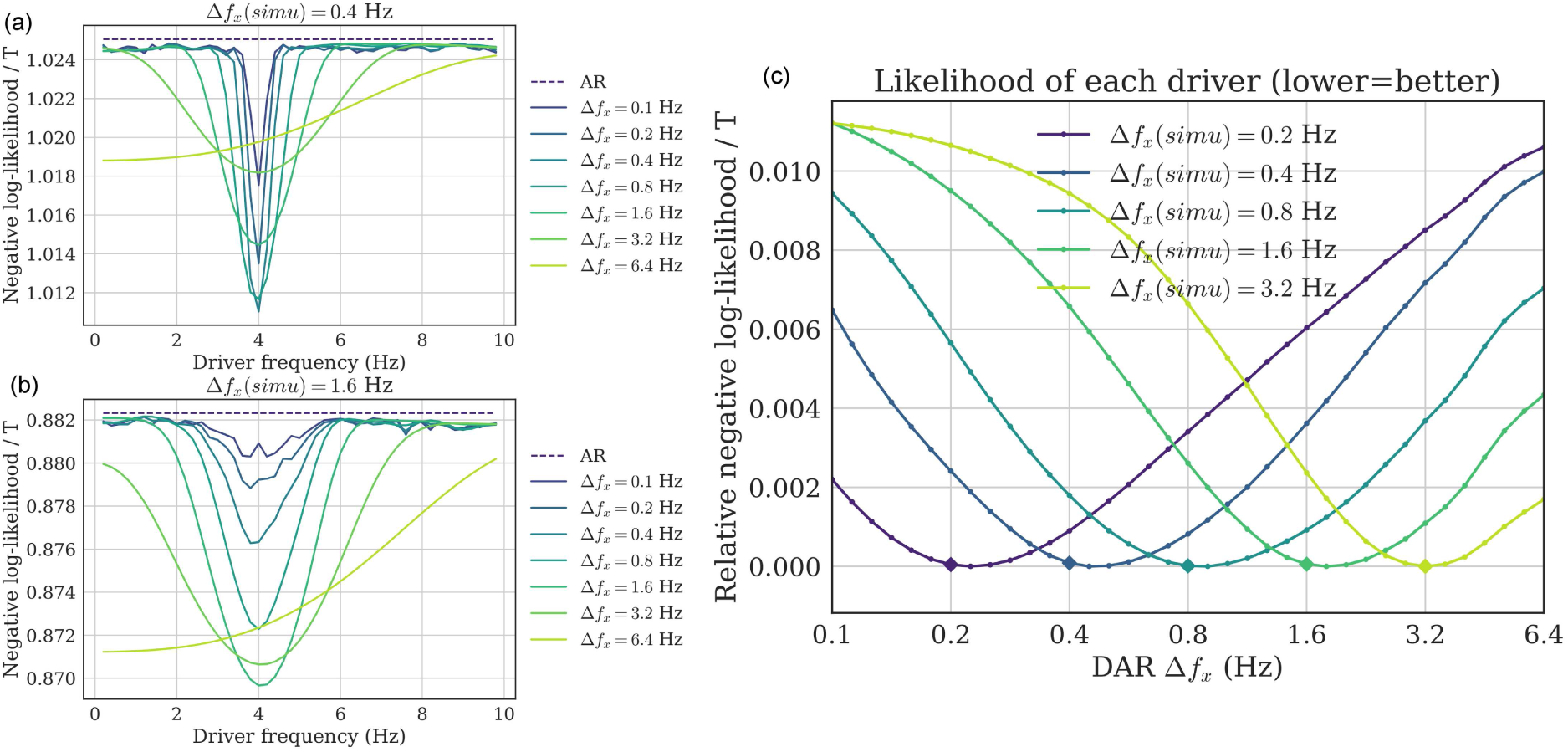
Driver’s frequency and bandwidth selection (simulations). (a,b) Two examples of grid-search over both *f*_*x*_ and ∆*f*_*x*_, over simulated signals with (a) ∆*f*_*x*_(*simu*) = 0.4 Hz and (b) *∆f*_*x*_(*simu*) = 1.6 Hz. For all bandwidths (except 6.4 Hz), the negative log-likelihood is minimal at the correct frequency *f*_*x*_ = 4 Hz. (c) To see more precisely the bandwidth estimation, we plot the negative log-likelihood (relative to each minimum for readability), for several ground-truth bandwidth ∆*f*_*x*_(*simu*), with *f*_*x*_ = 4 Hz. For each line, the minimum correctly estimates the ground-truth bandwidth (depicted as a diamond), showing empirically that the parameter selection method gives satisfying results.

This simulation confirms that the likelihood can be used to estimate the correct parameters for the driver’s filtering step, and that does not present any obvious bias in the estimation.

#### Driver estimates on human and rodent recordings

The outcome of the model selection procedure on the three real neurophysiological signals are reported in Fig. 5. Two general observations can be made. First, for all three signals, the optimal bandwidth was relatively large (3.2 Hz). Second, the optimal center frequency changed as we increased the bandwidth.

**Fig 5.**
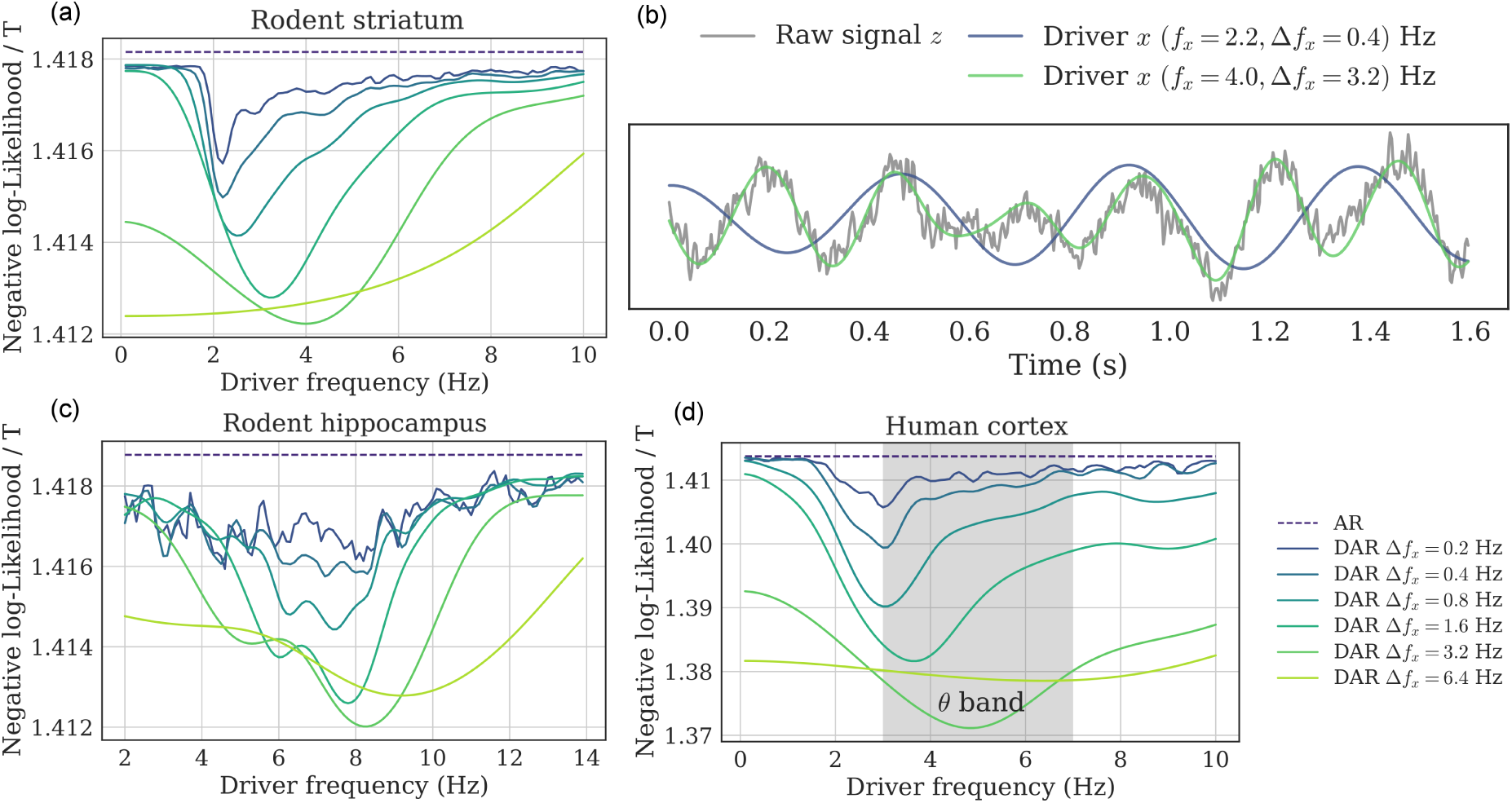
Driver’s frequency and bandwidth selection (real signals). (a,c,d) Negative log-likelihood of the fitted model for a grid of filtering parameters *f*_*x*_ and ∆*f*_*x*_: (a) rodent striatum, (c) rodent hippocampus, (d) human auditory cortex. The optimal bandwidth was very large (3.2 Hz), and the optimal center frequency changed as the bandwidth increased, suggesting that the optimal driver had a wide asymmetrical spectrum. (b) This portion of the rodent striatal signal shows two examples of driver with different bandwidths: The wide-band driver better follows the raw signal, and *independently* also leads to a better fit in DAR models.

Interestingly, this phenomenon was not observed in the simulation study, suggesting that in real data the optimal driver is wide-band and has an asymmetrical spectrum. In other words, the driver’s frequency is not precisely defined, and a large band-pass filter should be preferred to extract the driver.

These observations thus question which parameters should be used in practice. The classical approach is to build a comodulogram to select the best driver frequency *f*_*x*_, choosing arbitrarily the bandwidth ∆*f*_*x*_. This bandwidth is typically quite narrow, e.g. 0.4 Hz, and chosen to clearly isolate the maximum frequency in the comodulogram. However, this approach relies on the assumption that the driver is nearly sinusoidal, which is unrealistic [Cole and Voytek, 2017]. On the contrary, our data-driven approach selects the frequency and bandwidth that lead to the highest goodness of fit of our model.

To further describe the implications of our observations, we focused on the rodent striatal LFP signal. We compared an arbitrary choice of driver bandwidth (0.4 Hz) and the optimal choice with respect to the model likelihood (3.2 Hz). For these two bandwidths, we selected the optimal center frequencies, 2.2 Hz and 4.0 Hz respectively. Fig. 5(b) shows a time sample of the raw signal and the two extracted drivers. The wide-band driver (in green) followed very well the raw signal, which was not a perfect sinusoid. On the contrary, the narrow-band driver (in blue) seemed poorly related to the slow oscillation of the raw-signal. To know which of these two slow varying signals best captured the temporal amplitude modulations of the high frequencies, we fitted a model using each of these two drivers. We used the *same* high-pass filtered signal *y*, i.e. where we removed all frequencies below 16 Hz.

We found that the model with the wide-band driver explained more variance in the high frequencies than the model with the narrow-band driver, that is, that the model better explained the amplitude fluctuations of the high frequencies. As we used the same signal *y* for both models and as the driver lied in different frequency intervals, it should be noted that the fitting to the slow oscillation visible in Fig. 5(b) and the quality of the fit of the amplitude modulation in the high frequencies were two independent observations. By optimizing for the latter, one observe in Fig. 5(b) that it leads to a more realistic extraction of the driving neural oscillation.

More generally, our results emphasize the complexity of the choice of parameters in PAC analysis. With our method, we were able to easily compare different parameters, even on non-simulated data, which offered a principled way to set them and helped avoiding misinterpretation due to bad choices. We now investigate the conditional PSDs and the comodulograms estimates on these three datasets.

#### PSDs and comodulograms estimates of human and rodent neurophysiological recordings

Fig. 6 shows for each signal the PSD (in dB) depending on the driver’s phase, estimated through DAR models. The hyper-parameters (*p, m*) for the rodent striatal, the rodent hippocampal and the human cortical data were (90, 2), (15, 2), and (24, 2), respectively. These parameters were chosen by cross-validation with an exhaustive grid search: *p* ∈ [1, 100] and *m* ∈ [0, 3]. The filtering parameters (*f_x_, ∆f*_*x*_) were chosen to maximize the likelihood as described in previous section, and are respectively (8.2, 3.2) Hz, (5.2, 3.2) Hz and (4.0, 3.2) Hz (Fig. 6 bottom).

**Fig 6.**
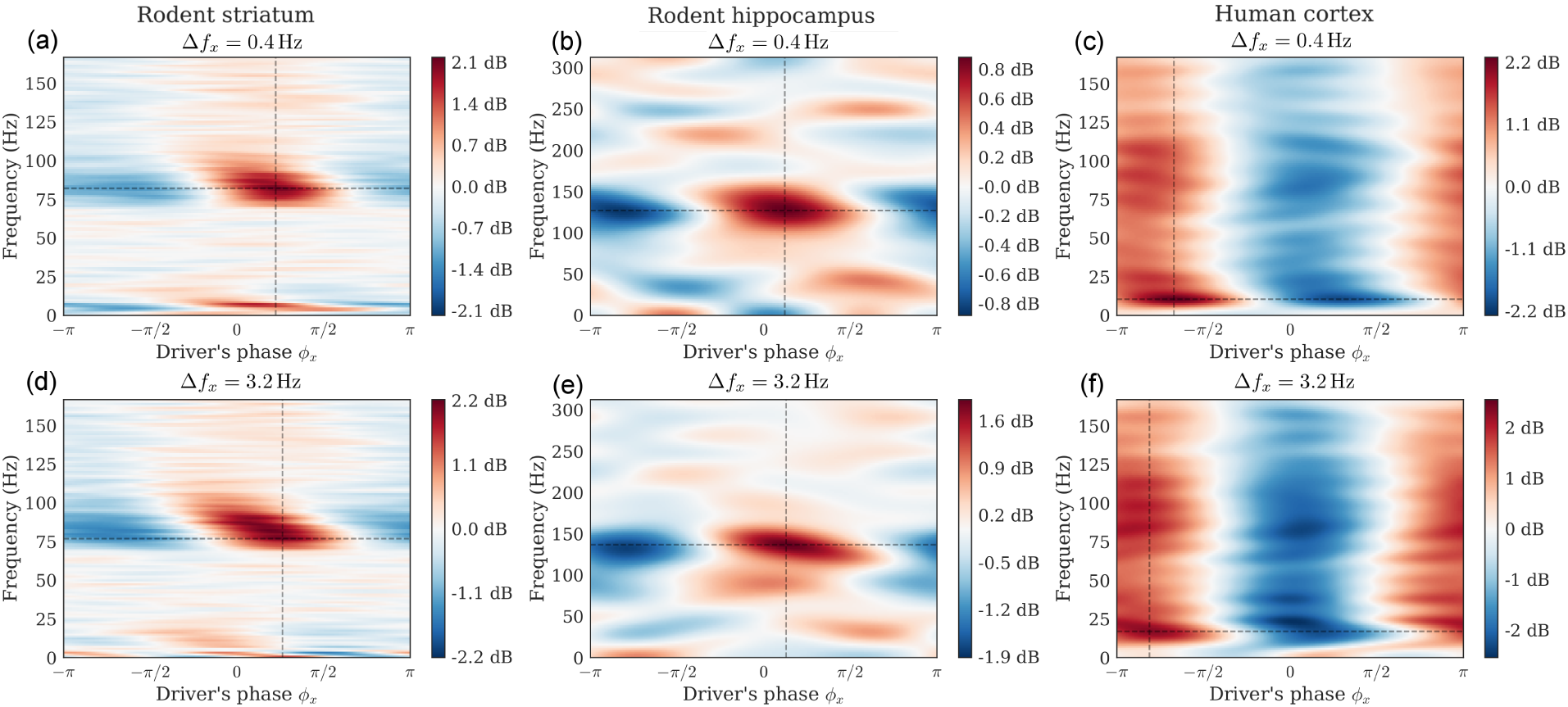
PSD conditional to the driver’s phase. Dataset: Rodent striatum (a,d), rodent hippocampus (b,e), human auditory cortex (c,f). We derive the conditional PSD from the fitted DAR models. In one plot, each line shows at a given frequency the amplitude modulation with respect to the driver’s phase. The driver bandwidth ∆*f*_*x*_ is 0.4 Hz (top row) and 3.2 Hz (bottom row). Note that the maximum amplitude is not always at a phase of 0 or π (i.e. respectively the peaks or the troughs of the slow oscillation). In figure (d), we can also observe that the peak frequency is slightly modulated by the phase of the driver (phase-frequency coupling).

For comparison, we also show the PSD obtained with a bandwidth ∆*f*_*x*_ = 0.4 Hz, with the center frequency chosen to maximize the likelihood. We used (6.4, 0.4) Hz, (3.2, 0.4) Hz and (2.2, 0.4) Hz (Fig. 6 top), respectively.

We observed two kinds of phenomena. In the rodent striatal and hippocampal data, the coupling was mainly concentrated around a given high frequency (i.e. 80 Hz and 125 Hz, respectively), whereas in the human cortical data, the coupling was observed at all frequencies, with a maximum around 20 Hz. The smoothness of the figures depends on the parameter *p*: a low value leads to a smoother PSD. Please note that the interpretation of the results depends on the filtering parameters.

The two phenomena can also be visualized in Fig. 7, where we plotted comodulograms for all three signals, computed with four different PAC metrics: the mean vector length first proposed in [Canolty et al., 2006] and updated by [Ӧzkurt and Schnitzler, 2011], the parametric model of [Penny et al., 2008] based on GLM, the modulation index of [Tort et al., 2010] that uses the Kullback-Leibler divergence, and our method based on DAR models.

**Fig 7.**
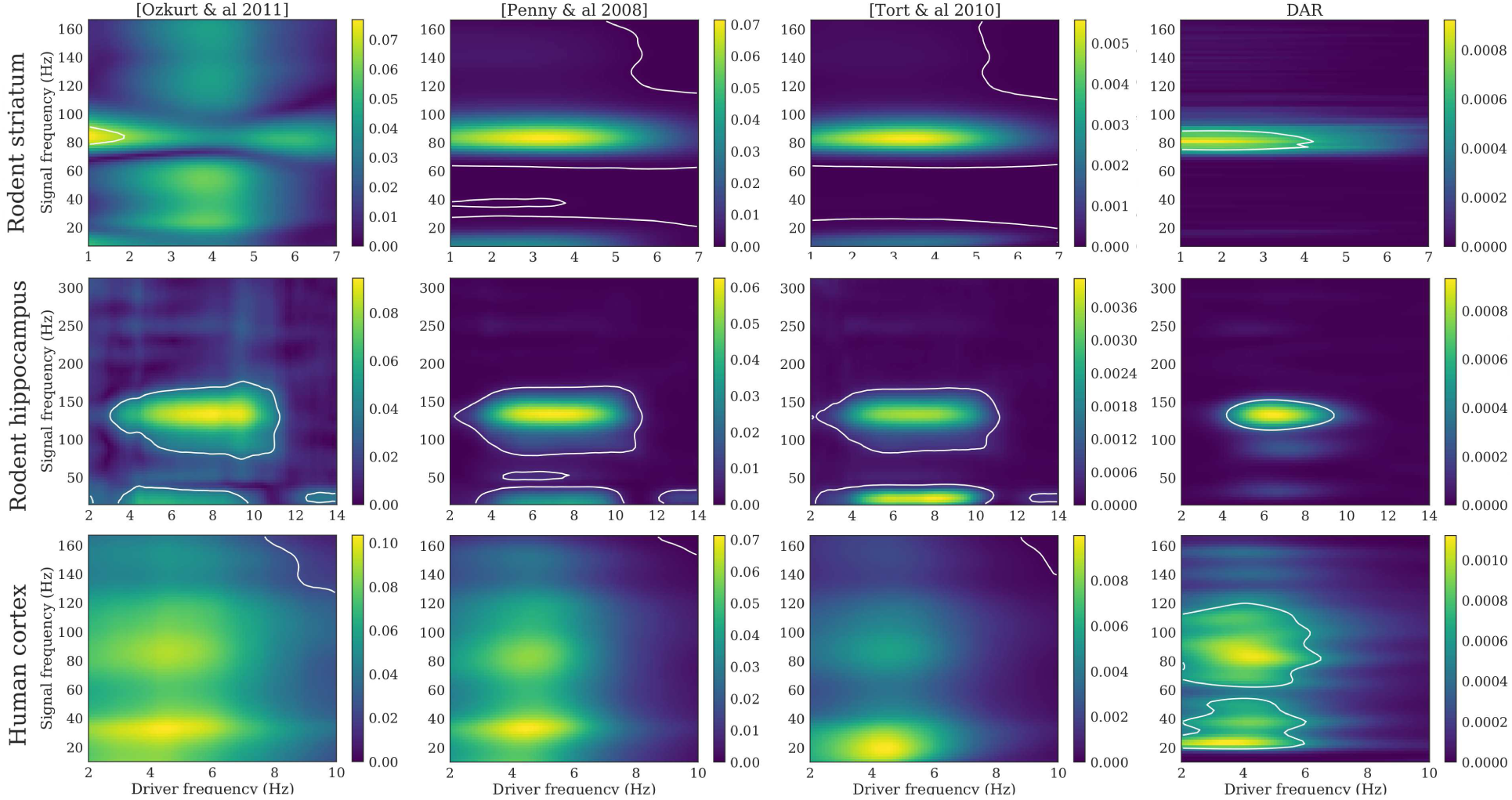
Comodulograms with parameters optimizing the likelihood. Dataset: rodent striatum (top), rodent hippocampus (middle) and human auditory cortex (down). Methods, from left to right: [Ozkurt *et al*. 2011] [Ӧzkurt and Schnitzler, 2011], [Penny *et al*. 2008] [Penny et al., 2008], [Tort *et al*. 2010] [Tort et al., 2010], DAR models (ours). The DAR model parameters and the driver bandwidth are chosen to be optimal with respect to the likelihood. White lines outline the regions with *p <* 0.01.

In DAR models, the hyper-parameters used are the cross-validated values described above. With other methods, high frequencies were extracted with a bandwidth ∆*f*_*y*_ twice the highest driver frequency used: 14 Hz, 28 Hz, and 20 Hz respectively. For all methods, the drivers were extracted with the optimal bandwidth *∆f*_*x*_ = 3.2 Hz.

The white lines crop the regions with a p-value *p <* 0.01. To compute such p-value, we added a random time shift between *x* and *y*, recomputed the entire comodulogram, and kept only the maximum value. We repeated this process 1000 times to have a distribution of maxima in the case of uncoupled surrogate signals.

Then, we took the 99-percentile of this distribution to obtain the threshold associated with the p-value *p* = 0.01. This method does not suffer from multiple testing issues contrary to the approach that estimates a different null distribution for each frequency couple in a comodulogram. We also avoided using a z-score since we cannot assume the distributions to be Gaussian.

The resulting comodulograms did not look like typically reported comodulograms, as we used a much larger bandwidth (3.2 Hz) than what was found in the literature. With an arbitrary choice of parameters, we could have obtained more classical comodulograms, as shown in Fig. 8. However, these results could be misleading because they suggest a coupling between sinusoidal oscillators, although the corresponding drivers are not perfectly sinusoidal (see Fig. 5(b)).

**Fig 8.**
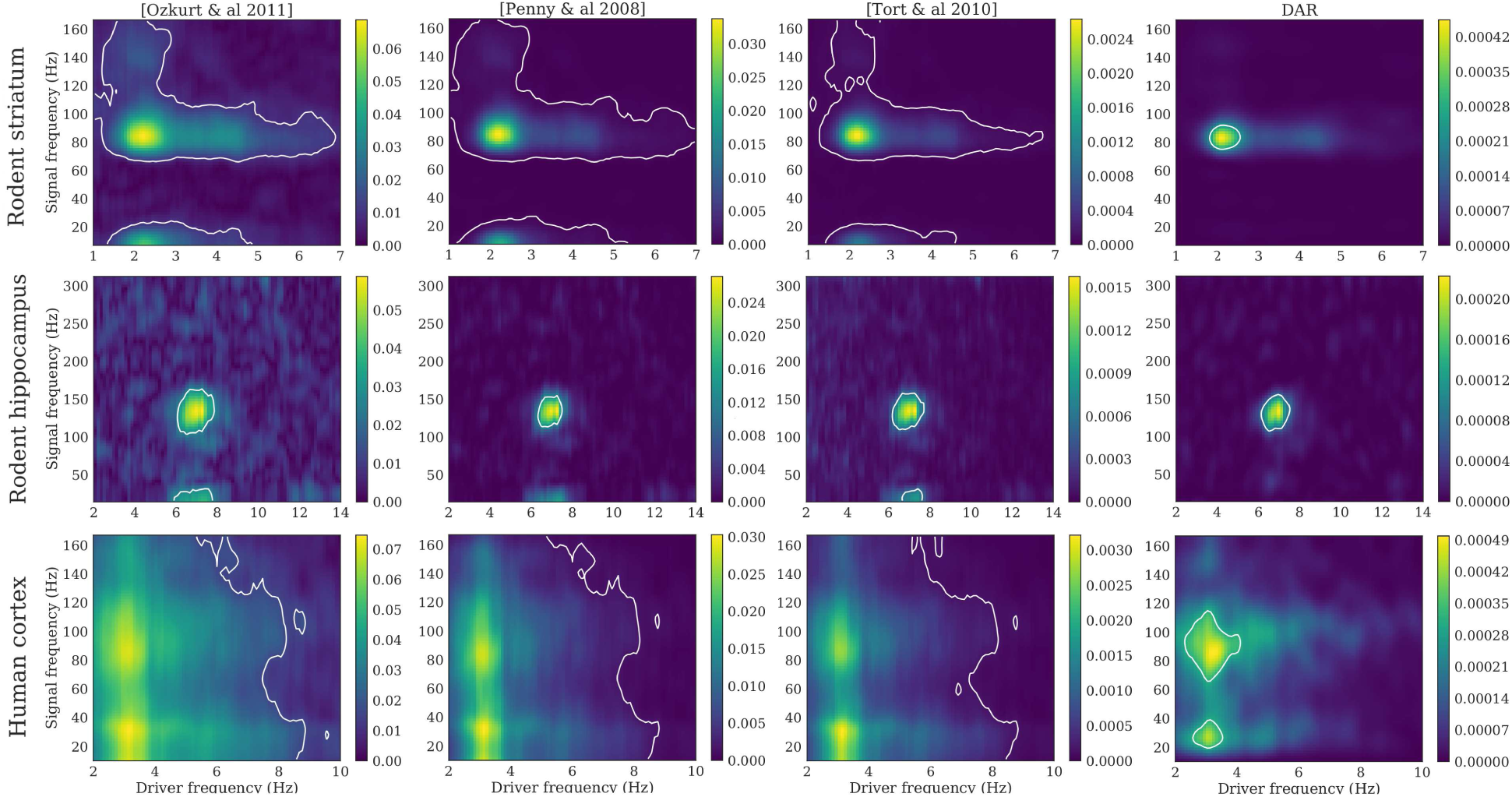
Comodulograms with parameters optimizing the comodulgoram sharpness. Dataset: rodent striatum (top), rodent hippocampus (middle) and human auditory cortex (down). Methods, from left to right: [Ozkurt *et al*. 2011] [Ӧzkurt and Schnitzler, 2011], [Penny *et al*. 2008] [Penny et al., 2008], [Tort *et al*. 2010] [Tort et al., 2010], DAR models (our). The driver bandwidth is chosen to have a well defined maximum in driver frequency: ∆*f*_*x*_ = 0.4 Hz. The DAR model parameters are chosen to give similar results than the other methods: (*p, m*) = (10, 2). White lines outline the regions with *p <* 0.01.

On these comodulograms, the four methods yielded comparable results as the data we used were very long: 1800, 100, and 730 seconds for the rodent striatal, the rodent hippocampal and the human cortical data, respectively. However, as we will see in the next section, differences emerge between methods when the signals are shorter.

#### Statistical power and robustness to small sample sizes

Given that DAR models are parametric with a limited number of parameters to estimate, less time samples may be needed to estimate PAC as compared to non-parametric methods. We tested this assumption using simulated signals of varying duration. We computed their comodulograms (as in Fig. 7) and selected the frequencies of maximum coupling. For each duration, we simulated 200 signals, and plotted the 2D histogram showing the fraction of time each frequency pairs corresponded to a maximum. We then compared the same four methods: DAR models with (*p, m*) = (10, 1), the GLM-based model [Penny et al., 2008], and two non-parametric methods [Tort et al., 2010, Ӧzkurt and Schnitzler, 2011]. Results shown in Fig. 9 show that parametric approaches provided a more robust estimation of PAC frequencies with short signals (*T* = 2 sec) than non-parametric methods.

**Fig 9.**
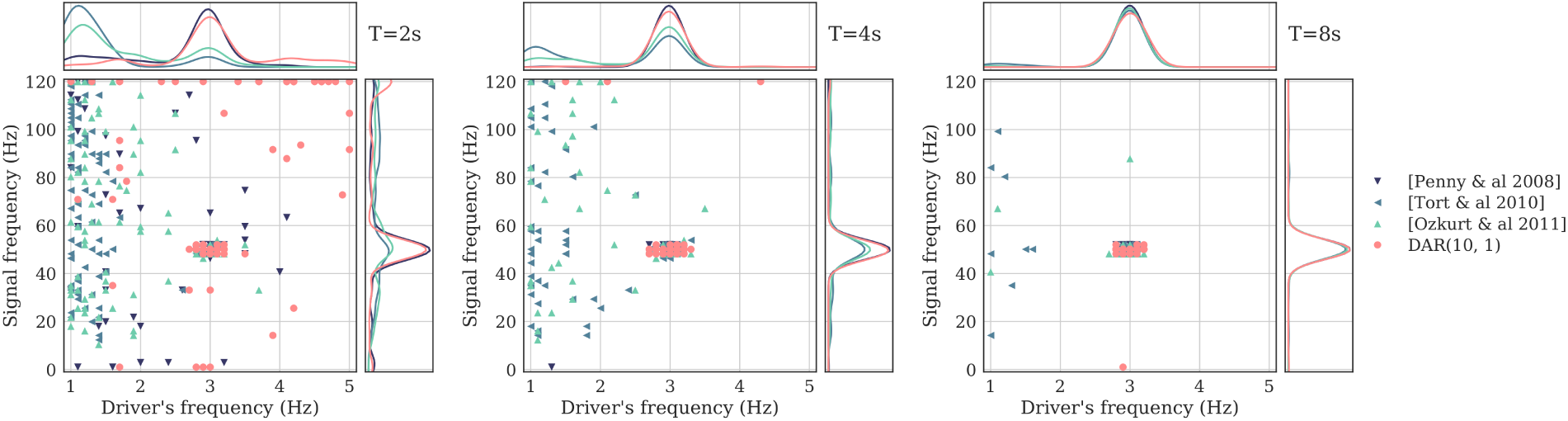
Robustness to small samples. Frequencies of the maximum PAC value, with four methods: a DAR model with (*p, m*) = (10, 1), the GLM-based model from Penny *et al*. [Penny et al., 2008], and two non-parametric models from Tort *et al*. [Tort et al., 2010] and Ozkurt *et al*. [Ӧzkurt and Schnitzler, 2011]. Each point corresponds to one signal out of 200. Kernel density estimates are represented above and at the right of each scatter plot. From left to right, the simulated signals last *T* = 2, 4 and 8 seconds. The signals are simulated with a PAC between 3 Hz and 50 Hz. The DAR models correctly estimate this pair of frequency even with a short signal length, as well as the GLM-based metric [Penny et al., 2008], while the two other metrics [Tort et al., 2010, Ӧzkurt and Schnitzler, 2011] are strongly affected by the small length of the signals, and do not estimate the correct pair of frequency.

The robustness to small sample size is a key feature of parametric models, as it significantly improves PAC analysis during shorter experiments. When undertaking a PAC analysis across time using a sliding time window, parametric models should therefore provide more robust PAC estimates. Note that the specific time values in these simulations should not be taken as general guidelines as they depend on the simulation parameters such as the signal-to-noise ratio. However, across all tests, parametric methods consistently provided more accurate results than non-parametric ones.

#### The amplitude fluctuations of the driver matter

One can note that in DAR models, the driver contains not only the phase of the slow oscillation, but also its amplitude. As the driver is not a perfect sinusoid, its amplitude fluctuates with time. On the contrary, most PAC metrics discard the amplitude fluctuations of the slow oscillation and only consider its phase. To evaluate these two options, we compared two drivers using DAR models: the original (complex) driver *x*(*t*), and the normalized driver 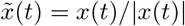. This normalized driver only contains the phase information, as in most traditional PAC metrics. Using cross-validation, we compared the log-likelihood of four fitted models, and found a difference always in favor of the non-normalized driver *x*(*t*), as it can be visualized in S3 Fig. This result shows that the coupling phenomenon is associated with amplitude fluctuations, a kind of phase/amplitude-amplitude coupling, as it was previously observed in [van Wijk et al., 2015]. Indeed, the GLM parametric method [Penny et al., 2008] was improved when taking into account the amplitude of the slow oscillation. Here, we use our generative model framework to provide an easy comparison tool through the likelihood, to validate this neuroscientific insight from the signals.

#### Directionality and delay estimation

In this section, we report the results of the directionality estimates using both simulations and neurophysiological signals. It is noteworthy that in DAR models, we arbitrarily call driver the slow oscillation although the model makes no assumption on the directionality of the coupling.

#### Simulations and comparisons

To validate the directionality estimations approach, we simulated signals with *T* = 1024 time points (4 seconds). We introduced a delay between the slow oscillation and the amplitude modulation of the fast oscillation, and verified that our method could correctly estimate the delays. We did not use a perfect sinusoidal driver, as the delay would only end up in a phase shift, and would not change the strength of the PAC. We used a bandwidth ∆*f*_*x*_ = 2 and a noise level σ_*ε*_ = 0.4.

We compared our method, using a DAR model with (*p, m*) = (10, 1), to the cross-frequency directionality (CFD) approach described in [Jiang et al., 2015]. This method makes use of the phase slope index (PSI) [Nolte et al., 2008] for PAC estimation. In [Jiang et al., 2015], the PSI over a frequency band *F* is defined as:

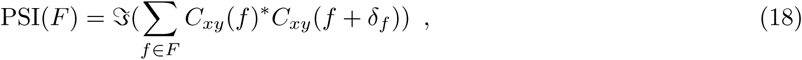

where *C*_*xy*_ is the complex coherence, 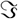 is the imaginary part, and δ_*f*_ is the frequency resolution. As the PSI was not designed to provide an estimation of the delay, we *modify* it into:

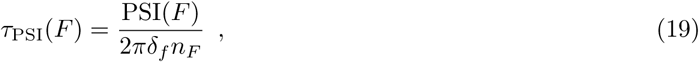

where *n*_*F*_ is the number of frequencies in set *F*. Note that this delay estimator was correct only if the coherence was almost perfect *C*_*xy*_(*f*) ≈ 1 and if the phase slope was small enough to have sin(****ϕ****(*f* + δ_*f*_) − ****ϕ****(*f*)) ≈ ****ϕ****(*f* + δ_*f*_) − ****ϕ****(*f*). As these assumptions are rarely met in practice, we only used this estimator to compare *qualitatively* with our own estimator.

Results presented in Fig. 10 show the mean estimated delay with ±1 standard deviation computed over 20 simulations. The delay was correctly estimated in both time directions, with no visible bias. As expected, the modified CFD was biased, and showed a higher variance than the DAR-based approach.

**Fig 10.**
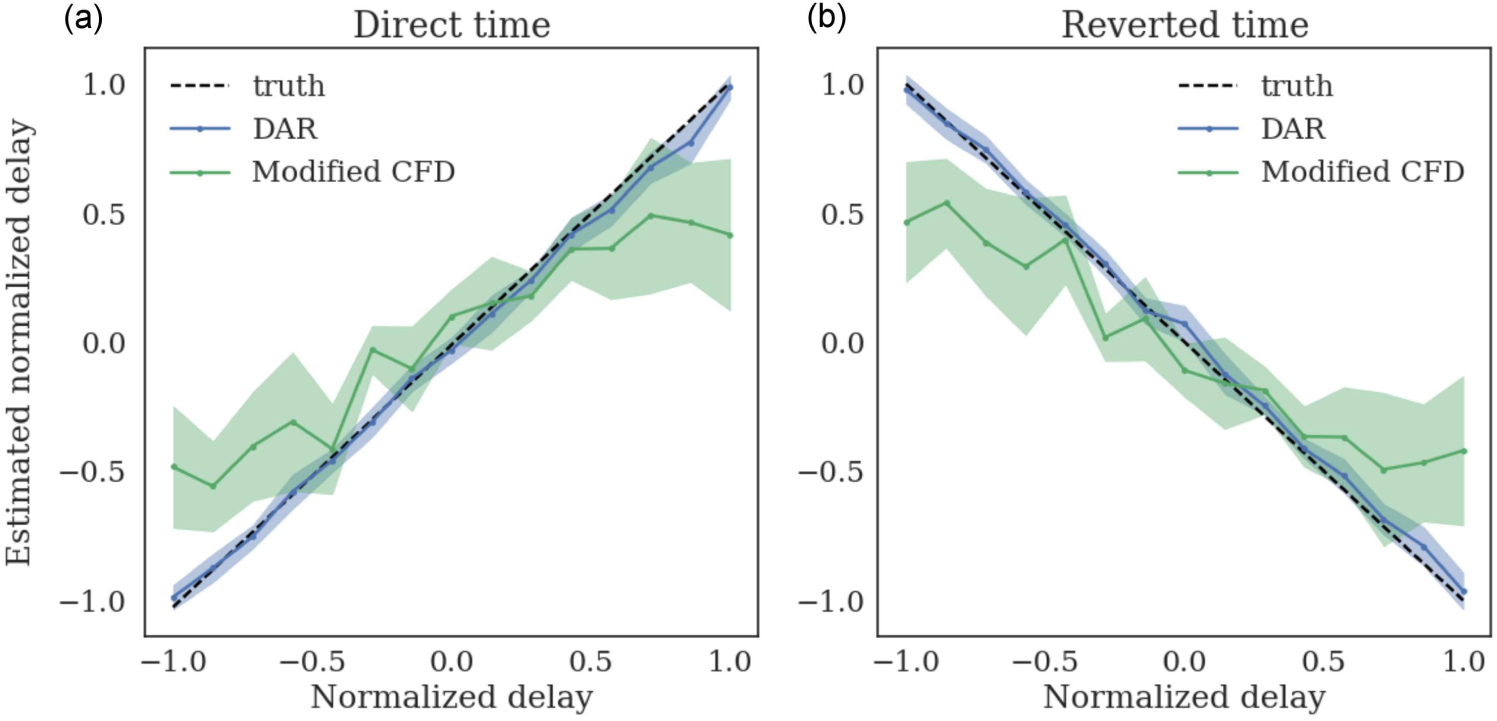
Estimating delays over 20 simulations for each delay. The delays are estimated on both direct time (a) and reverted time (b). The delays are normalized: τ = 1 corresponds to one driver oscillation, 1*/f*_*x*_ sec. The modified CFD shows a bias and only serves for qualitative comparison. The delay estimation based on DAR models correctly estimates the delays, with no visible bias.

#### Delay estimates of human and rodent data

We applied this delay estimation method on the three experimental recordings, with both direct time and reverted time. Once again, note that the comparison with the CFD [Jiang et al., 2015] is only qualitative, since the CFD was not designed as a delay estimator. The temporal delay was estimated with DAR models with the parameters obtained from the cross-validation selection. The driver we used was obtained with the best filtering parameters found in our grid-search: (*f_x_, ∆f*_*x*_) are respectively (4.0, 3.2) Hz, (8.2, 3.2) Hz, and (5.2, 3.2) Hz.

The results are presented in Fig. 11, where we display the delay estimated with maximum likelihood over multiple DAR models. We also display error bars which correspond to the standard deviation obtained with a block-bootstrap strategy [Carlstein, 1986]. This strategy consists in splitting the signal into *n* = 100 non-overlapping blocks of equal length, to draw at random (with replacement) *n* blocks, and to evaluate the delay on this new signal. We repeated this process 20 times and computed the standard deviation of the 20 estimated delays. Such strategy is used to estimate empirically the variance of a general statistic from stationary time series.

**Fig 11.**
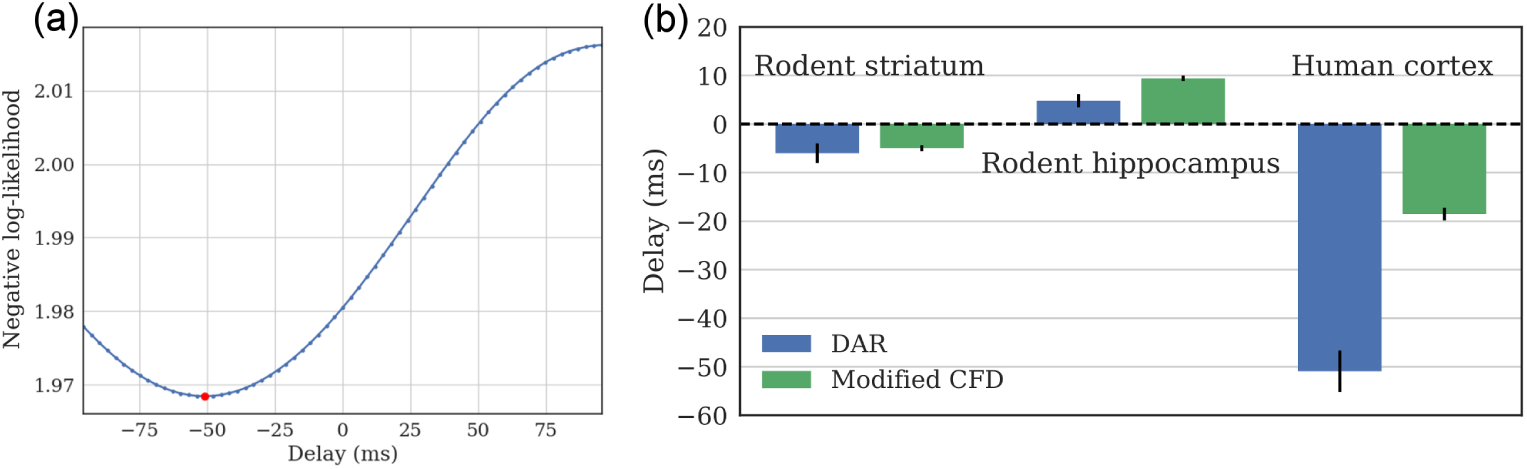
Estimated delays on rodent and human datasets. (a) Negative log likelihood for multiple delays, with the human cortical signal. (b) Optimal delay for the three signals, computed with model selection on DAR models, and with a CFD method modified to provide a delay. The error bars indicate the standard deviation obtained with a block-bootstrap strategy. For the rodent hippocampal data, the delay is positive: the low frequency oscillation precedes the high frequency oscillation. For the human cortical data and the rodent striatal data, the delay is negative: The high frequency oscillation precedes the low frequency oscillation.

First, one can observe that the results were qualitatively similar between the two methods. The CFD was biased toward zero for large delays, as was previously shown in the simulations. One can also note that the directionality was not always the same. For the rodent striatal and the human cortical data, the delay was negative and the best model fit happened between the signal *y* and the *future* driver *x*. Inversely, in the rodent hippocampal data, the delay was positive, and the best model fit was between the signal *y* and the *past* driver *x*. Further experiments need to be performed to better understand the origin of such delays but we demonstrate here the use of DAR models to estimate them.

Note that our method does not select the delay that leads to the maximum PAC, which would not be statistically valid. On the contrary, we select the delay that leads to the best fit of the model on the data. This approach is more rigorous since we maximize the variance explained by our model, and not the effect of interest.

## Discussion

Cross-frequency coupling (CFC) and phase-amplitude coupling (PAC) more specifically have been proposed to play a fundamental role in neural processes ranging from the encoding, maintenance and retrieval of information [Buzsáki, 2010, Jensen and Colgin, 2007, Lisman and Jensen, 2013, Axmacher et al., 2010, Fell and Axmacher, 2011, Kaplan et al., 2014, Hyafil et al., 2015], to large-scale communication across neural ensembles [Canolty and Knight, 2010, Jirsa and Müller, 2013, Khan et al., 2013, Florin and Baillet, 2015].

While a steady increase in observations of PAC in neural data has been seen, how to best detect and quantify such phenomena remains difficult to settle. We argue that a method using DAR models, as described here, is rich enough to capture the time-varying statistics of brain signals in addition to provide efficient inference algorithms. These non-linear statistical models are probabilistic, allowing the estimation of their goodness of fit to the data, and allowing for an easy and fully controlled comparison across models and parameters. In other words, they offer a unique principled data-driven model selection approach, an estimation strategy of phase/amplitude-amplitude coupling based on the approximation of the actual signals, a better temporal resolution of dynamic PAC and the estimation of coupling directionality.

One of the main features of PAC estimation through our method is the ability to compare models or parameters on non-synthetic data. On the contrary, traditional PAC metrics *cannot* be compared on non-synthetic data, and two different choices of parameters can lead to different interpretations. There is no legitimate way to decide which parameter shall be used with empirical data using traditional metrics. The likelihood of the DAR model that can be estimated on left-out data offers a rigorous solution to this problem.

We presented here results on both simulated signals and empirical neurophysiological signals. The simulations gave us an illustration of the phenomenon we want to model, and helped us understand how to visualize a fitted DAR model. They also served a validation purpose for the bandwidth selection approach that we performed on real data. Using the data-driven parameter selection on non-synthetic signals, we showed how to choose sensible parameters for the filtering of the slow oscillation. All empirical signals are different, and it was for example reported in the neuroscience literature that peak frequencies vary between individuals [Haegens et al., 2014] and that this should not be overlooked in the analysis of the data. The parameter selection based on fitted DAR models makes it possible to fit parameters on individual datasets. Our results also shed light on the asymmetrical and wide-band properties of the slow oscillation, which could denote crucial features involved in cognition [Cole and Voytek, 2017].

The second novelty of our method stands in considering the amplitude fluctuations of the slow oscillation in the PAC measure and not only its phase. Using the rodent and human data, we showed that the instantaneous amplitude of the slow oscillation influences the coupling in PAC, as it was previously suggested in [van Wijk et al., 2015]. The amplitude information should therefore not be discarded as it is done by existing PAC metrics. For instance, the measure of alpha/gamma coupling reported during rest [Osipova et al., 2008, Roux et al., 2013] should incorporate alpha fluctuations when studied in the context of visual tasks [Voytek et al., 2010], as an increase of alpha power is often concomitant with a decrease of gamma power [Fries et al., 2001]. The comparison between DAR models considering or not these low-frequency power fluctuations would inform on the nature of the coupling: purely phase-amplitude, or rather phase/amplitude-amplitude. In Tort *et al*. [Tort et al., 2008], both theta power changes and modulation of theta/gamma PAC were reported in rats having to make a left or right decision to find a reward in a maze.

The use of our method could decipher whether the changes in coupling were related to the changes in power, informing on the underlying mechanisms of decision-making. Moreover, as our method models the entire spectrum simultaneously, a phase-frequency coupling could potentially be captured in our models. Therefore, our method is not limited to purely phase-amplitude coupling, and extends the traditional CFC analysis.

Furthermore, in those types of experiments, changes in PAC can be very fast depending on the cognitive state of the subject. Therefore, the need for dynamic PAC estimates is growing [Tort et al., 2008]. We showed with simulations that DAR models are more robust than non-parametric methods when estimating PAC on small time samples. This robustness is critical for time-limited experiments and also when analyzing PAC across time in a fine manner, typically when dynamic processes are at play.

Last but not least, likelihood comparison can also be used to estimate the delay between the coupled components, which would give new insights on highly debated questions on the role of oscillations in neuronal communication [Fries, 2005, Bastos et al., 2015]. For example, a delay close to zero could suggest that the low and high frequency components of the coupling might be generated in the same area, whereas a large delay would suggest they might come from different areas. As an alternative interpretation, the two components may come from the same area, but the coupling mechanism itself might be lagged. In this case, a negative delay would suggest that the low frequency oscillation is driven by the high frequency oscillations, whereas a positive delay would suggest that the low frequency oscillation drives the high frequency amplitude modulation. In any case, this type of analysis will provide valuable information to guide further experimental questions.

A recent concern in PAC analysis is that all PAC metrics may detect a coupling even though the signal is not composed of two cross-frequency coupled oscillators [Kramer et al., 2008, Lozano-Soldevilla et al., 2016, Amiri et al., 2016, Gerber et al., 2016, Vaz et al., 2017]. It may happen for instance with sharp slow oscillations, described in humans intracranial recordings [Cole et al., 2016]. Sharp edges are known not to be well described by a Fourier analysis, which decomposes the signal in a linear combination of sinusoids. Indeed, such sharp slow oscillations create artificial high frequency activity at each sharp edge, and these high frequencies are thus artificially coupled with the slow oscillations. This false positive detection is commonly referred to as “spurious” coupling [Jensen et al., 2016]. Fig. 12 shows a comodulogram computed on a simulated spurious PAC dataset, using a spike train at 10 Hz, as described in [Gerber et al., 2016]. The figure shows that all four methods, including the proposed one, detect some significant PAC, even though there is no nested oscillations in the signal. Even though our method does not use filtering in the high frequencies, it does not solve this issue and is affected in the same way as other traditional PAC metrics. Indeed, our work shed light on the wide-band property of the slow oscillations, but DAR models cannot cope with full-band slow oscillations, which contain strong harmonic components in the high frequencies. However, we consider that such “spurious” PAC can also be a relevant feature of a signal, as stated in [Cole et al., 2016]. In their study, they show that abnormal beta oscillations (13-30 Hz) in the basal ganglia and motor cortex underlie some “spurious” PAC, but are actually a strong feature associated with Parkinson’s disease. A robust way to disentangle the different mechanisms that lead to similar PAC results remains to be developed.

**Fig 12.**
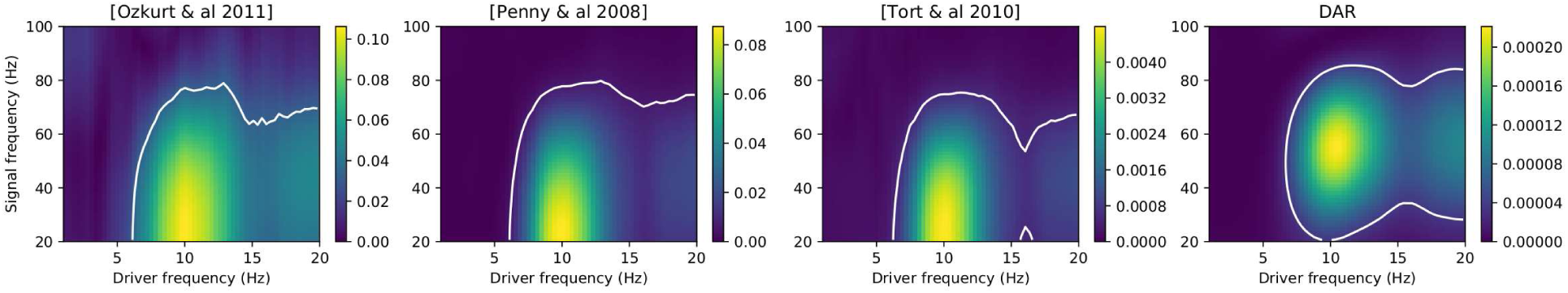
Comodulograms obtained on a simulated signal with spurious PAC. Spurious PAC was generated using a spike train at 10 Hz, as described in [Gerber et al., 2016]. All four methods, including the proposed one, detect some significant PAC, even though there is no nested oscillations.

The method we presented in this paper uses univariate signals obtained invasively in rodents or humans.

As a lot of neurophysiological research uses non-invasive MEG or EEG recordings containing multiple channels, a multivariate analysis could be of high interest. One way to use data from multiple channels is to estimate a single signal using a spatial filter such as in [Cohen, 2017]. Such a method is therefore complementary to univariate PAC metrics like ours which can be applied to the output of the spatial filter. The method from [Cohen, 2017] builds spatial filters that maximize the difference between, say, high-frequency activity that appears during peaks of a low-frequency oscillation *versus* high-frequency activity that is unrelated to the low-frequency oscillation. Again, from the signal obtained with the spatial filter, it is straightforward to adapt most PAC metrics such as our method.

Neurophysiological signals have all the statistical properties to make them a challenge from a signal processing perspective. They contain non-linearities, non-stationarities, they are noisy and they can be long, hence posing important computational challenges. Our method based on DAR models offer novel and more robust possibilities to analyse neurophysiological signals, paving the way for new insights on how our brain functions via spectral interactions using local or distant coupling mechanisms.

Inline with the open science philosophy of this journal, our method is fully available as an open source package that comes with documentation, tests, and examples: https://pactools.github.io

## Acknowledgments

We would like to thank Khodagholy *et al*. [Khodagholy et al., 2015] for sharing rodent hippocampal LFP sample data and Canolty *et al*. [Canolty et al., 2006] for sharing human cortical ECoG sample data. We also thank two anonymous reviewers for their excellent feedback and suggestions on a previous version of our manuscript.

This work was supported by the ERC Starting Grant SLAB ERC-YStG-676943 to Alexandre Gramfort, the ERC Starting Grant MindTime ERC-YStG-263584 to Virginie van Wassenhove, the ANR-16-CE37-0004-04 AutoTime to Valérie Doyère and Virginie van Wassenhove, and the IDEX NoTime to Valérie Doyère, Alexandre Gramfort and Virginie van Wassenhove,.

## Supporting Information

**S1 Fig.**
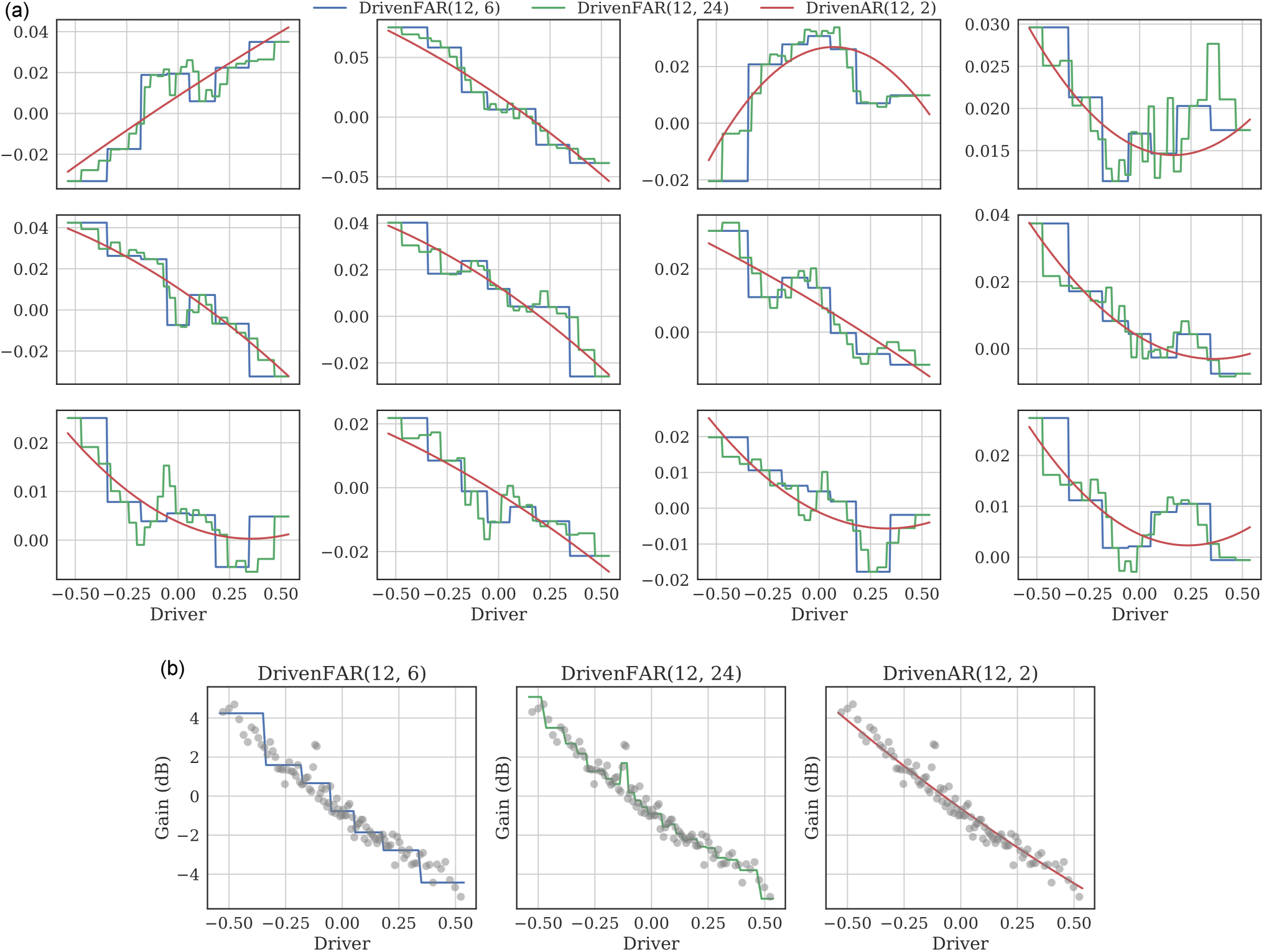
Validating the polynomial basis for DAR. AR coefficients *a*_*i*_(*x*) as functions of the driver *x*. (b) Innovation variance σ^2^(*x*) as functions of the driver *x*. The gray circles are the mean squared-residual over 100 bins of the driver values. We compared three parametrizations of these functions: Two staircase functions with respectively *s* = 7 and *s* = 25 steps, and one polynomial function with order *m* = 2, as used in the rest of this work. For a piecewise constant AR parametrization with *s*-steps, we divided the driver’s values into *s* equally distributed bins, then we fitted *s* independent linear AR models on the time samples of each bin. This approach is more general than the polynomial approach, yet each AR model uses only *T /s* samples, whereas the polynomial approach in the DAR model uses all the *T* samples. The DAR model is therefore more robust. The polynomial parametrization also uses fewer coefficients since a low order *m* is sufficient. The models are fitted on the human cortical signal, using *p* = 12. We see that an order-2 polynomial is sufficient to approximate the trend of both the AR coefficients and the innovation variance, as estimated by the staircase functions, yet the polynomial uses much fewer parameters.

**S2 Fig.**
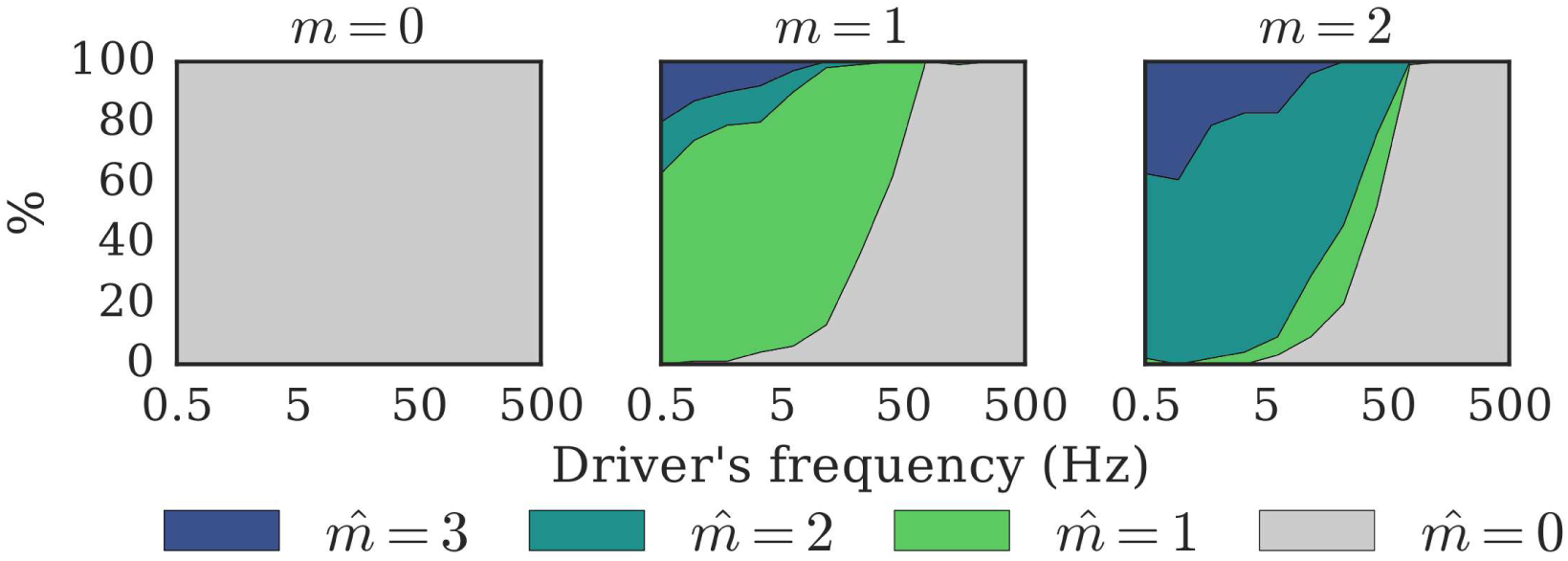
Model selection with the Bayesian information criterion (BIC). We simulated 100 signals from 100 DAR models with *p* = 10 and *m* ∈ [0, 1, 2]. We then estimated new DAR models on these signals, with *p* ranging from 1 to 20, and *m* ranging from 0 to 3. We selected 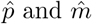 that minimized the BIC. The graphs show the proportion of runs that lead to each value of 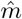. The hyper-parameter *m* is correctly estimated in most cases if the driver’s frequency is not too high (*f*_x_ < 50 Hz). The hyper-parameter *p* is correctly estimated at ±2 in most cases.

**S3 Fig.**
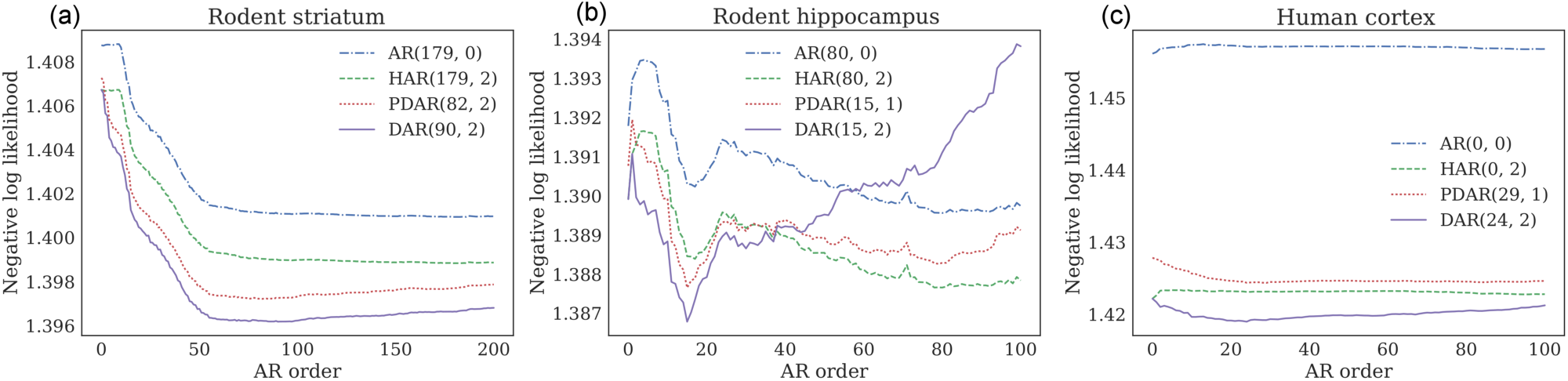
Model selection with cross-validation. Cross-validation on our three dataset, (a) rodent striatum, (b) rodent hippocampus, and (c) human auditory cortex, to select the best model and the best parameters. Splitting the signal in half, we fitted the models on the first half, and evaluated the model likelihood on the second half. We compared four different models on a grid of parameter *p* ∈ [0, 100 − 200] and *m* ∈ [0, 3]: 1. AR: a linear AR model 2. Heteroskedastic AR (HAR): an hybrid model between a linear AR model and a DAR model, where the innovation variance σ^2^ is driven by *x*, but the AR coefficients are constant in time. 3. Phase DAR (PDAR): a DAR model, with a normalized driver: *x/|x|*. In this way, we only consider the phase of the slow oscillation, as in most PAC metrics. 4. DAR: a DAR model, where both the innovation variance σ^2^ and the AR coefficients are driven by *x*. The figures present the negative log likelihood (lower is better) by time sample. Each line corresponds to a given model with its best parameter *m*. The legend shows which orders (*p, m*) are the best for each model. One can observe that the curves of negative log-likelihood are not convex, yet they exhibit rather clear minima used to define the optimal paramaters.

**S4 Fig.**
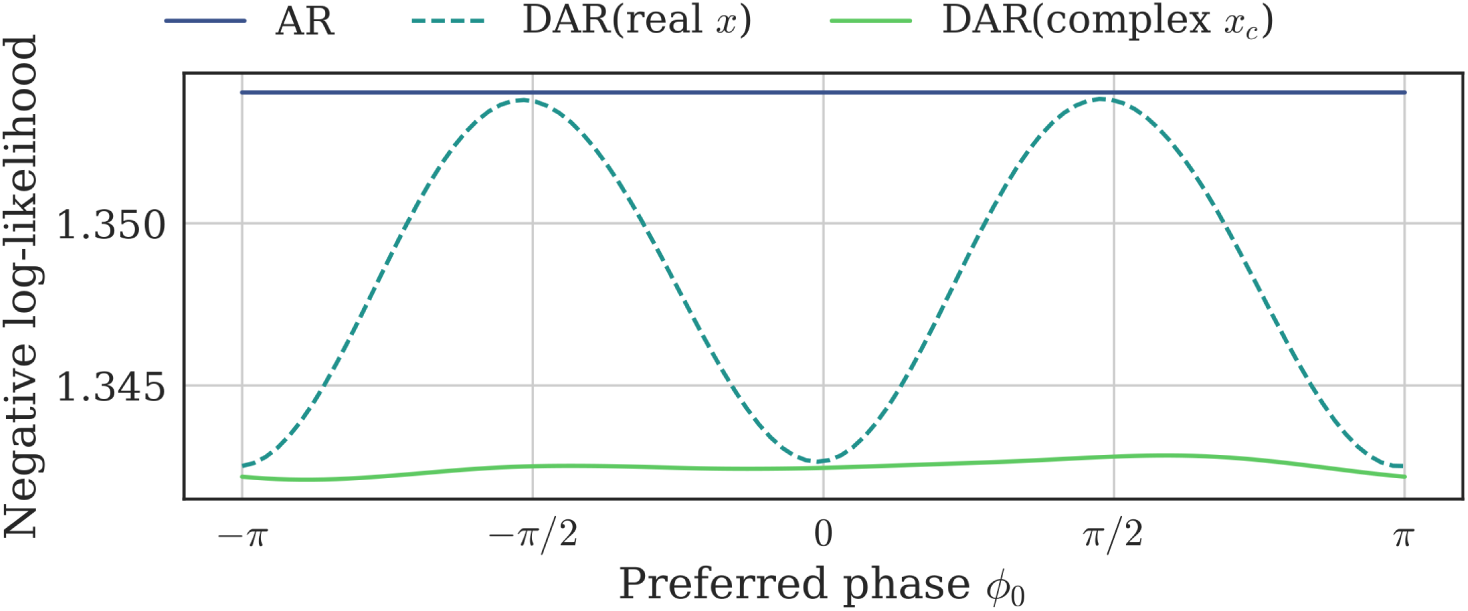
Removing the bias with a complex-valued driver. We simulated a signal as described in the Methods section, introducing a phase difference ****ϕ****_0_ in the modulation. For each value of ****ϕ****_0_, we fitted three different models, and compared their negative log-likelihood by time sample (the lower the better): an AR, a DAR with a real driver, and a DAR with a complex driver. The parameters were set to *p* = 10 and *m* = 1. A bias is visible around ****ϕ****_0_ = ±π*/*2, since the real-valued driver DAR model does not fit better than the AR model. As expected, this bias disappears when we update the model to a complex-valued driver DAR model.

